# Cavin1 binding nanobodies reveal structural flexibility and regulated interactions of the N-terminal coiled-coil domain

**DOI:** 10.1101/2024.11.26.625551

**Authors:** Ya Gao, Vikas A. Tillu, Yeping Wu, James Rae, Thomas E. Hall, Kai-en Chen, Saroja Weeratunga, Qian Guo, Emma Livingstone, Wai-Hong Tham, Robert G. Parton, Brett M. Collins

## Abstract

Caveolae are abundant plasma membrane structures that regulate signalling, membrane homeostasis, and mechanoprotection. Their formation is driven by caveolins and cavins and their coordinated interactions with lipids. We have developed nanobodies against the trimeric HR1 coiled-coil domain of Cavin1. We identify specific nanobodies that do not perturb Cavin1 membrane binding and localise to caveolae when expressed in cells. The crystal structure of a nanobody HR1 complex reveals a symmetric 3:3 architecture as validated by mutagenesis. In this structure, the C-terminal half of the HR1 domain is disordered suggesting the nanobody has stabilised an open conformation previously identified as important for membrane interactions. A phosphomimic mutation in a Thr-Ser pair proximal to this region reveals selective regulation of Cavin2/Cavin3 association. These studies provide new insights into Cavin domains required for assembly of multiprotein caveolar assemblies and describe new nanobody tools for structural and functional studies of caveolae.

**SUMMARY STATEMENT:** Nanobodies are reported that can label the Cavin1 protein and caveola structures in cells.

## INTRODUCTION

The nanoscale (∼70 nm) cell-surface invaginations of the plasma membrane called caveolae (‘little caves’) are abundant in many vertebrate cells including endothelial cells, cardiac muscle fibers and adipocytes (Lamaze et al., 2017; Parton, 2018; Parton et al., 2020b). Caveolae possess a distinct protein and lipid composition consistent with their roles in physiological processes including endocytosis (Boucrot et al., 2011), regulation of intracellular signaling (Couet et al., 1997; Garcia-Cardena et al., 1997), lipid homeostasis (Harder and Simons, 1997; Kenworthy et al., 2023; Zhou et al., 2021), and mechanoprotection (Dewulf et al., 2019; Gervasio et al., 2011; Lo et al., 2015; Lolo et al., 2022; Lundmark et al., 2024; Sens and Turner, 2006; Sinha et al., 2011). Caveola dysfunction or loss has been implicated in a range of human diseases including cardiovascular arrhythmia (Campostrini et al., 2017; Fridolfsson and Patel, 2013; Jia and Sowers, 2015; Vatta et al., 2006), lipodystrophy and muscular dystrophy (Hayashi et al., 2009; Parton, 2018; Parton et al., 2020a; Rajab et al., 2010), highlighting their importance in animal physiology. Lipids and proteins act together to shape the plasma membrane associated caveolae, which requires the assembly of two distinct protein families, caveolins and cavins, and their coordinated interactions with membrane lipids and cholesterol (Kenworthy et al., 2023; Ludwig et al., 2013; Lundmark et al., 2024; Parton et al., 2020b; Rothberg et al., 1992; Tillu et al., 2021; Zhou et al., 2021).

The integral membrane caveolins (CAV1, CAV2 and muscle specific CAV3) are synthesized as 8S-CAV oligomers at the endoplasmic reticulum which are then trafficked to the plasma membrane via the Golgi apparatus forming higher order oligomeric assemblies (Busija et al., 2017; Hayer et al., 2010; Morales-Paytuvi et al., 2023). CAV1 is sufficient to generate membrane invaginations in bacterial model systems that have the same size as typical caveolae in mammalian cells (Walser et al., 2012), and is organized as a homo-oligomer of 11 CAV1 subunits in a spiraling flat disc that sits in the inner leaflet of the plasma membrane (Porta et al., 2022). However, in mammalian cells CAV1 requires the presence of cavins to generate morphological caveolae (Hansen et al., 2009; Hayer et al., 2010; Hill et al., 2008; Liu and Pilch, 2008). The cavins are peripheral membrane proteins that form a multiprotein complex constitutively assembled in the cytosol and subsequently associated with 8S-CAV1 complexes at the plasma membrane (Bastiani et al., 2009; Hill et al., 2008). There are four different isoforms of cavin proteins in mammalian cells, Cavin1-4, which exhibit distinctive tissue-specific roles (Parton et al., 2020a). Cavin1 is ubiquitously expressed and is essential for stabilizing the caveolar domain and promoting the formation of the invaginated caveola structure (Hill et al., 2008). Cavin2 and Cavin3 are in competition with one another for interacting with Cavin1 to form higher order hetero-oligomeric complexes that regulate the size and function of the caveola coat (Bastiani et al., 2009; Gambin et al., 2013; Ludwig et al., 2013; McMahon et al., 2019; Mohan et al., 2015).

All cavin proteins share a common primary structure possessing two highly conserved positively-charged regions, helical region 1 (HR1) and helical region 2 (HR2), separated by three poorly conserved disordered regions (DR1, DR2 and DR3) along their length (Burgener et al., 1990; Gustincich and Schneider, 1993; Hill et al., 2008; Kovtun et al., 2014; Tillu et al., 2018). The highly basic HR domains and uniformly acidic DR domains presumably produces a highly polarised cavin protein molecule (Bastiani et al., 2009). The molecular mechanism of caveola biogenesis is rapidly emerging from structural studies (Lundmark et al., 2024), including cryoEM structures of CAV1 (Han et al., 2024; Porta et al., 2022) and crystal structures of Cavin1 and Cavin4 HR1 domains (Kovtun et al., 2014). The cavin HR1 domains form a highly extended trimeric coiled-coil assembly and provides a core structure thought to mediate intermolecular interactions of cavins promoting homo- and hetero-oligomerization. A positively-charged basic surface towards the C-terminus of the HR1 domain interacts preferentially with plasma membrane-enriched phosphatidylinositol 4,5-bisphosphate (PtdIns(4,5)P_2_) for lipid-dependent recruitment of cavins to caveolar membranes and membrane remodelling activity (Hansen et al., 2009; Kovtun et al., 2014; Zhou et al., 2021). Notably, this region of the HR1 domain is also found to undergo local unfolding during membrane interaction, leading to partial insertion of hydrophobic residues (Liu et al., 2022). The C-terminal HR2 domain of Cavin1 contains an eleven-residue repeated amino acid sequence, termed the undecad repeat (UC1) that further promotes efficient binding with phosphatidylserine (PtdSer) for membrane remodelling (Kovtun et al., 2014; Tillu et al., 2018). The HR2 domain is also functionally important for the self-assembly of cavins into larger oligomers and together with HR1 associates with anionic lipid membranes (Kovtun et al., 2014). The three intrinsically unstructured DR domains of Cavin1 are strictly required for formation of caveola invagination and dynamic trafficking of caveolae (Tillu et al., 2021). Therefore, the assembly of caveolae has been proposed to depend on the low affinity electrostatic interaction between CAV1 and Cavin1 as well as membrane lipid interactions (Kenworthy et al., 2023; Tillu et al., 2021).

With the aim of developing novel tools and reagents to study cavin structure and function we have turned to nanobodies. These single domain antibodies derived from camelid species (e.g. alpacas and llamas) have allowed the development of new therapeutics and research tools for selective detection of proteins in their native environment, with advantages over antibodies or small molecules due to their high affinity, stability, improved solubility and yield (Jin et al., 2023). Using the mouse Cavin1 HR1 complex we immunised alpacas and subsequently isolated two high affinity nanobodies, NbA12 and NbB7, that specifically target this Cavin1 coiled-coil domain. Crystal structures and mutagenesis define their binding epitopes within the N-terminal half of the HR1 domain. Interestingly, in the crystal structure of the NbB7-Cavin1 HR1 complex the C-terminal region of the HR1 domain is disordered, suggesting the nanobody is bound to a conformation that may resemble the proposed semi-unfolded state induced upon membrane-binding. We further show that while NbA12 does not associate with caveolae in cells, NbB7 is robustly recruited to endogenous caveolae in a Cavin1-dependent manner. The unfolding of Cavin1 detected with NbB7 occurs at a central hydrogen-bonded pair of residues within the coiled-coil sequence. Mutagenesis of this region shows that it is critical for specificity between Cavin2 and Cavin3 recruitment in cells. These nanobodies thus appear to be recognising epitopes that are differentially accessible *in vitro* and *in vivo*, and can be used as selective tools for structural analyses and cellular imaging of caveola assemblies.

## RESULTS

### Identification of highly potent nanobodies targeting Cavin1

Nanobodies against the folded trimeric coiled-coil HR1 domain of mouse Cavin1 (mC1-HR1; residues 45-155) were produced by immunization of an alpaca with purified HR1 domain. Using a nanobody phage display library, we identified nanobodies that bind conformational epitopes of properly folded HR1 domain. From 43 positive phage supernatants that recognize mouse Cavin1 HR1 domain by preliminary ELISA screening, we identified 7 distinct clonal groups based on unique CDR3 sequences, and expressed and purified one representative from each clonal group for further characterisation. Of these nanobodies NbA12 and NbB7 were found to show the strongest interaction with mC1-HR1 domain by GST-pulldown assay, with a potential weaker interaction with NbA4 (**Figure 1A**). These were then validated by isothermal titration calorimetry (ITC), and NbA12 and NbB7 were both found to bind purified mC1-HR1 with nanomolar affinities (**Figure 1B; Table S1**). In contrast NbA4 did not show any significant binding affinity by ITC (**Figure S1A**). Sequences of NbA12 and NbB7 are provided in **Figure S1B**.

**Figure 1.**
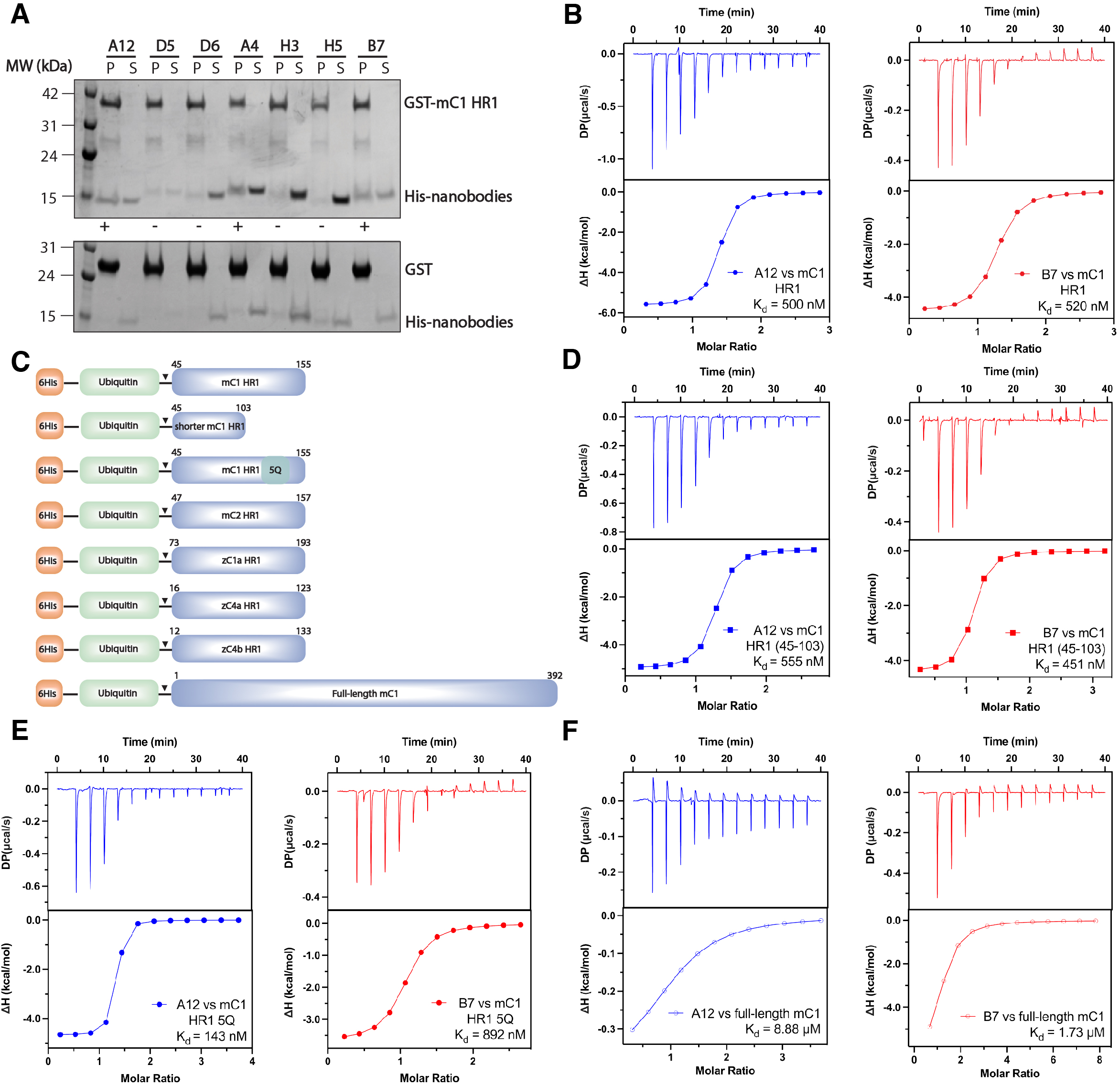
Identification of Cavin1 binding nanobodies. **(A)** Initial selections of mouse Cavin1 HR1 domain with nanobodies. GST-mC1 HR1 was used as a bait for His-nanobodies. **(B)** ITC thermogram for binding of NbA12 and NbB7 to mC1-HR1. **(C)** Schematic representation of HR1 domain and mutations in cavin family proteins tested for nanobody binding (shown to scale). **(D)** NbA12 and NbB7 show identical binding affinity with an N-terminal fragment of mC1-HR1(45-103) domain measured by ITC. **(E)** Similarly, both NbA12 and NbB7 interact with the 5Q mutant of mC1-HR1 domain measured by ITC. **(F)** Binding NbA12 and NbB7 with full-length mouse Cavin1 protein was still detected by ITC, although the affinity is reduced compared to the isolated HR1 domain. For all ITC graphs, the upper panel represents raw data, and the lower panel represents the normalized and integrated binding isotherms fitted with a 1-to-1 binding ratio. The binding affinity (K_d_) is determined by calculating the mean of at least two independent experiments. Thermodynamic parameters for the binding analysis are provided in **Table S1.**

To map the binding regions of the nanobodies we tested a series of mC1-HR1 truncation and mutant constructs (**Figure 1C**). An N-terminal fragment mC1-HR1(45-103) retains the stable trimeric coiled-coil conformation when expressed and purified from *E. coli* (Kovtun et al., 2014) and displays an almost identical binding affinity for both NbA12 and NbB7 nanobodies (**Figure 1D**) as the full HR1 domain (**Figure 1B and 1D**). This indicates that C-terminal region of HR1 domain is not required for binding with these nanobodies. Previously, substitutions of several key charged Arg and Lys residues (Lys115, Arg117, Lys118, Lys124, Arg127) with Gln residues located in a basic cluster at the C-terminus of the Cavin1 HR1 domain (5Q mutant) were made that disrupt its association with negatively-charged phospholipid membranes but still maintain proper helical structure (Kovtun et al., 2014). Consistent with binding of the N-terminal region of the HR1 domain, we found that the binding affinities of the 5Q mutant for nanobody NbA12 or NbB7 were not significantly different to the wild-type HR1 domain (**Figure 1E**).

Next, we tested binding of NbA12 to mC1-HR1 in the presence of a molar excess of NbB7 and found that binding was no longer detectable (**Figure S1C**). This suggests that they are most likely sharing an overlapping epitope on the HR1 coiled-coil structure. Lastly, we tested these two nanobodies against full-length MBP-tagged mCavin1. Notably, although both nanobodies still showed significant interactions, they were found to have reduced binding affinities compared to the isolated HR1 domain alone (**Figure 1F; Table S1**). The full-length mCavin1 protein is relatively difficult to purify compared to the HR1 domain alone, but we previously showed it can also form larger molecular weight oligomers of the core Cavin1 trimer (Kovtun et al., 2014; Tillu et al., 2021). We speculate that potential intra- and/or intermolecular interactions of the HR1 domain with other regions of the Cavin1 protein may be partially occluding the nanobody binding site.

Given the high affinity of both nanobodies for the mouse Cavin1 HR1 domain immunogen, we then sought to test the interaction of nanobodies with HR1 domain in other cavin proteins to examine its binding specificity. NbA12 and NbB7 both showed binding with mouse Cavin2 HR1 (mC2-HR1) but with much lower affinity (*K_d_*) in the range of 3.4-7.7 μM (**Figure S1D**). Next, we also tested the binding of the nanobodies with cavin homologues from zebrafish to see if they showed cross-species reactivity. A similar micromolar binding affinity result was detected where we observed binding between both nanobodies and Cavin1a HR1 domain from zebrafish (zC1a-HR1) (**Figure S1E**). Zebrafish Cavin4a HR1 (zC4a-HR1) has been confirmed to form a similar trimeric coiled-coil structure by X-ray crystallography (Kovtun et al., 2014). In contrast to nanobody NbB7, which showed a weak binding in the micromolar range, nanobody NbA12 was unable to bind to zebrafish Cavin4a HR1 (**Figure S1F**). For zebrafish Cavin4b, these two nanobodies showed no interaction (**Figure S1G**). In summary, these two nanobodies are relatively specific for mouse Cavin1, but possess some weak cross reactivity with mouse Cavin2 and with cavin homologues in other species.

### Cavin1 retains lipid binding and remodelling activity in the presence of nanobodies

The Cavin1 HR1 domain was previously shown to interact with negatively charged phospholipid membranes promoting recruitment of cavins to caveolar membranes and membrane remodelling activity *in vitro* (Kovtun et al., 2014). We first tested whether nanobodies affect membrane binding by the HR1 domain with artificial liposomes in pelleting assays. For this experiment we used liposomes composed of bovine brain-derived Folch fraction which is highly enriched in negatively charged phospholipids. As shown previously (Kovtun et al., 2014), using GST-tagged mC1-HR1 we confirmed a high affinity association with Folch liposomes (**Figure 2A**). This interaction was not affected by either NbA12 or NbB7, which were both co-pelleted with the GST-tagged mC1-HR1. Full-length Cavin1 was also shown to have an ability to remodel and tubulate artificial liposomes *in vitro* (Kovtun et al., 2014). As shown in (**Figure 2B**) the nanobodies did not perturb the ability of full-length recombinant Cavin1 to remodel and tubulate artificial liposomes *in vitro*. Altogether these results indicate that the binding of the nanobodies does not inhibit the key membrane interacting activities of Cavin1.

**Figure 2.**
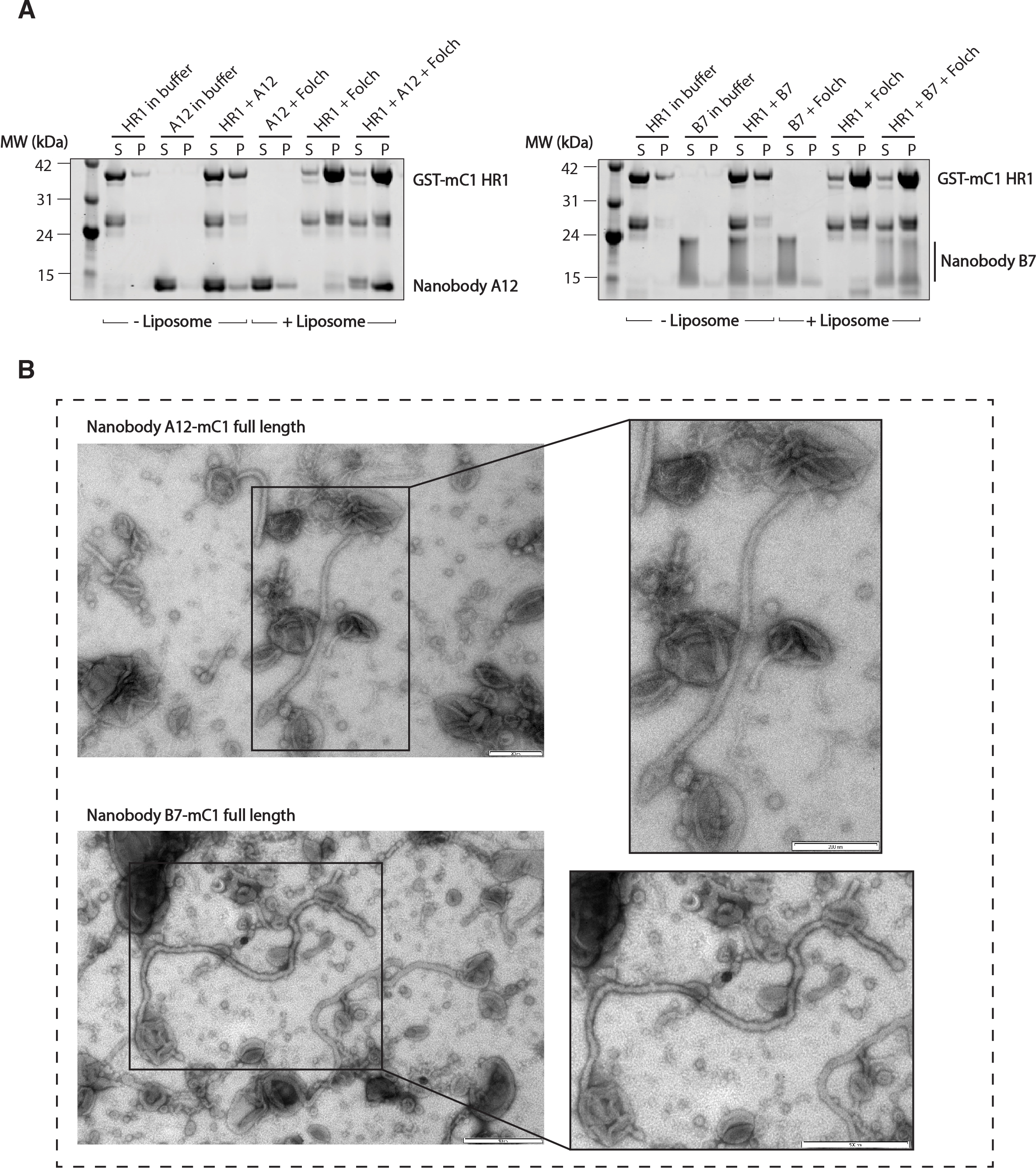
Nanobodies do not affect Cavin1’s *in vitro* intrinsic membrane remodelling properties. **(A)** Liposome pelleting assay of mouse Cavin1 HR1 domain in the presence of nanobodies. Multilamellar vesicles were generated from Folch I lipids. In the presence or absence of NbA12 or NbB7, mouse Cavin1 HR1 domain was incubated with or without liposomes and centrifuged, “S” and “P” stand for unbound supernatant and bound pellet respectively. **(B)** Negatively stained electron microscopy images of nanobody-Cavin1 complexes incubated with Folch I liposomes. His-MBP full-length mouse Cavin1 was used to test liposome tubulation in the presence of either nanobody NbA12 or NbB7. Scale bars of Cavin1-nanobody NbB7 = 500 nM. Scale bars of Cavin1-nanobody NbA12 = 400 nM.

### NbB7 labels Cavin1-coated caveolae in cells

As both nanobodies NbA12 and NbB7 can bind Cavin1 *in vitro* and do not perturb its ability to associate with membranes, we next tested their capacity to label Cavin1 in several different cell lines including HeLa and A431 human lines and baby hamster kidney (BHK) cells. Their interaction with the two structural proteins of caveolae, Cavin1 and CAV1, were firstly validated by immunoprecipitation in transiently transfected HeLa cells. Lysates from cells expressing C-terminal GFP-tagged NbA12-GFP and NbB7 were incubated with GFP-binding nanobody (Kubala et al., 2010) covalently coupled to NHS-activated Sepharose^™^ resin. Under these co-immunoprecipitation conditions with mild detergent (0.5% Triton X-100) only NbB7-GFP could co-precipitate endogenous Cavin1, and to a lesser extent CAV1 (**Figure 3A**). One important note is that, for unknown reasons, GFP-tagged NbB7 and NbA12 nanobodies migrate at different molecular weights by SDS-PAGE. The experiment has been repeated several times with similar results, and plasmids were re-sequenced to confirm that there were no differences at the nucleotide levels for these constructs. In A431 cells with lower expression levels we could detect NbB7-GFP at the correct molecular weight with an additional band at the smaller size, suggesting that some truncation or nicking of the protein may be occurring (**Figure S1H**).

**Figure 3.**
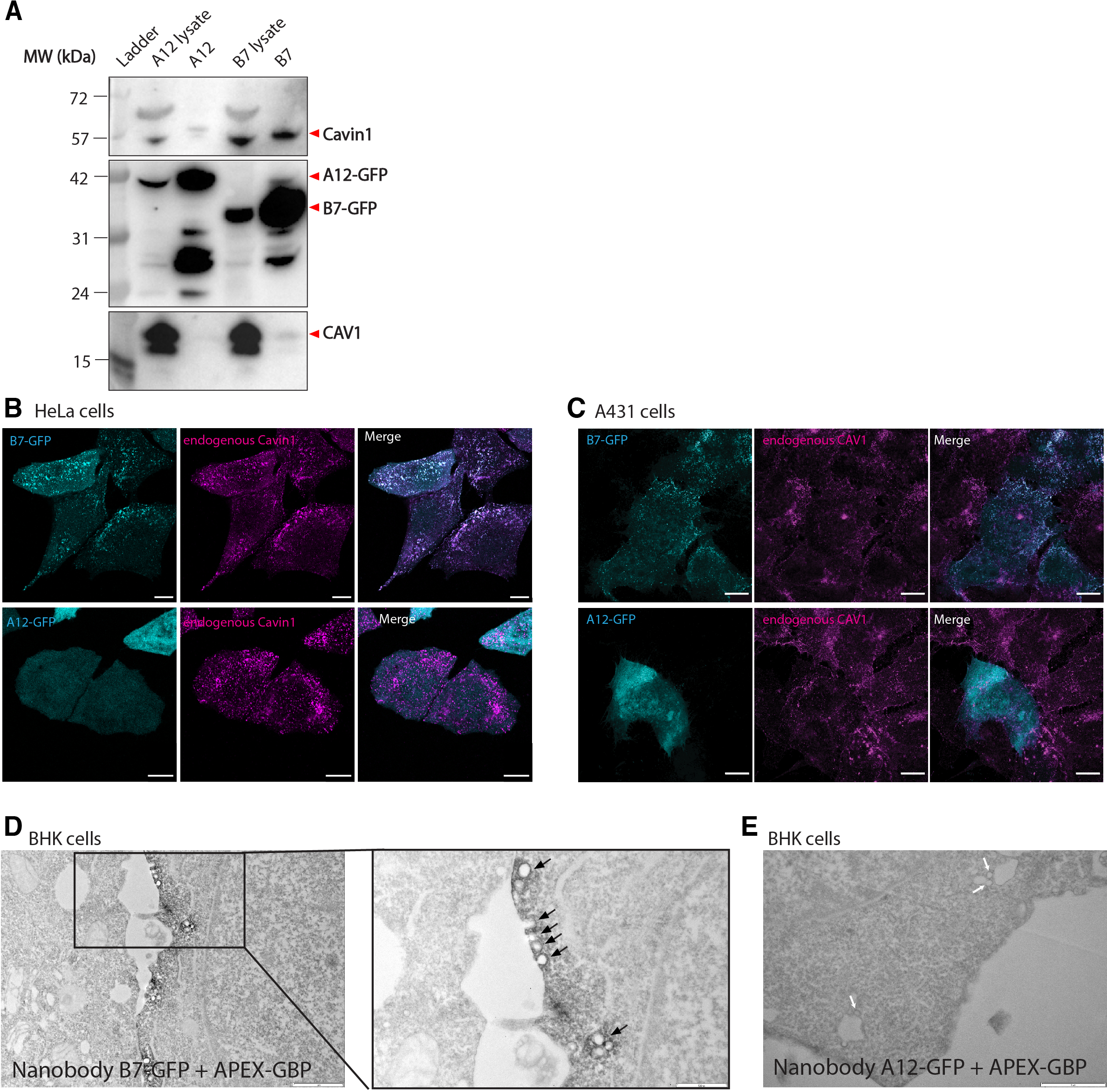
NbB7 can be used as a molecular probe for Cavin1 in cells. **(A)** Co-immunoprecipitation of GFP-nanobodies with Cavin1 and CAV1. HeLa cells expressing NbA12-GFP, NbB7-GFP were incubated with GFP-nanobody couple NHS-activated Sepharose^™^ resin. Cavin1 and CAV1 were detected by Western blot. **(B)** Confocal microscopy of HeLa cells transfected with NbB7-GFP (upper) and NbA12-GFP (bottom) nanobody. Fixed cells were also stained for Cavin1. Scale bars = 10 μM. **(C)** Confocal microscopy of A431 cells transfected with NbB7-GFP (upper) and NbA12-GFP (bottom) nanobody. Fixed cells were also stained for CAV1. Scale bars = 10 μM. **(D)** NbB7-GFP nanobody detected at cell plasma membrane by APEX-GBP co-transfection in BHK cells. Scale bars = 1 μM (left) and 500 nM (right expanded view). Black arrows represent invaginated caveolae labelled with APEX-GBP. (**E**) NbA12-GFP nanobody showed cytosolic labelling by APEX-GBP co-transfection (right) in BHK cells. White arrows indicate caveolae without APEX-GBP labelling. Scale bar = 1 μM.

We next investigated the co-localization of nanobodies with Cavin1 and CAV1 using an orthogonal set of cells by fluorescence microscopy. HeLa cells and A431 cells express both endogenous Cavin1 and CAV1 and form typical caveola puncta at the cell plasma membrane. Consistent with the co-immunoprecipitation results we observed a high degree of colocalization between NbB7-GFP and endogenous Cavin1 at the plasma membrane of HeLa cells, while in contrast NbA12-GFP was diffusely expressed in the cytoplasm (**Figure 3B**). NbB7-GFP nanobody also showed a high degree of co-localization with endogenous CAV1 in A431 cells with a characteristic punctate caveola distribution, while NbA12-GFP nanobody again showed no co-localization with CAV1 (**Figure 3C**). The lack of Cavin1/caveola association of NbA12 is consistent with its poor binding in immunoprecipitation experiments and suggests that although NbA12 can bind Cavin1 *in vitro,* its required binding epitope may not be accessible in the membrane assembled caveola complex. As NbB7-GFP shows clear association with caveolae by fluorescence imaging we next examined its localisation by electron microscopy. The APEX tag, derived from soybean ascorbate peroxidase, has been developed as a fast, sensitive, and reliable labelling system by fusing it to a GFP-binding peptide (GBP) that allows direct imaging of target proteins fused to GFP tag (Ariotti et al., 2015). Here, we co-transfected baby hamster kidney (BHK) cells with NbB7-GFP and APEX-GBP in cells. NbB7-GFP labelling with APEX-GBP revealed an electron-dense reaction product strictly associated with caveolae at the cell plasma membrane (**Figure 3D**). The co-expression of GFP-nanobody NbA12 and APEX-GBP resulted in a cytosolic labelling with no visible caveolae labelled (**Figure 3E**). This high-resolution protein localization analysis indicates that nanobody NbB7 is highly suitable for use as a bioactive tool to explore cavins and caveolae in cells.

### NbB7 binding promotes partial unfolding of the HR1 coiled-coil structure

We next aimed to characterize the molecular basis of nanobody binding to cavin1. We co-crystallized the mC1-HR1 trimeric coiled-coil domain with NbA12 and NbB7. Attempts to obtain diffraction quality crystals with NbA12 were unsuccessful, however a stable stoichiometric complex of nanobody NbB7– mC1-HR1 purified by size exclusion chromatography was crystallised by sparse-matrix screening and optimization. The structure of the nanobody NbB7–mC1-HR1 complex was solved by X-ray crystallography at 1.5 Å resolution. This revealed a symmetric complex with three copies of NbB7 bound to the mC1-HR1 trimer, with trimeric symmetry along the crystallography 3-fold axis. The structure showed clear electron density for nanobody NbB7 interacting with the N-terminal region of the trimeric coiled-coil HR1 residues 56-98 (of residues 45-155 in the crystallised construct) (**Figure 4A, 4B & 4C; Table S2**). One notable feature of the structure, however, is that when the Cavin1 HR1 protein forms a complex with nanobody NbB7, the C-terminal half of the HR1 coiled-coil appears to have unravelled, with no visible electron density for HR1 residues 99-155. It is clear that if this region formed an extended coiled-coil as previously observed for apo-mouse Cavin1 and zebrafish Cavin4 HR1 domain structures (Kovtun et al., 2014) it would not be compatible with the packing of the crystal lattice. Unfolding of the C-terminus of the HR1 domain was recently proposed to be important for membrane interaction (Liu et al., 2022) and is discussed further below.

**Figure 4.**
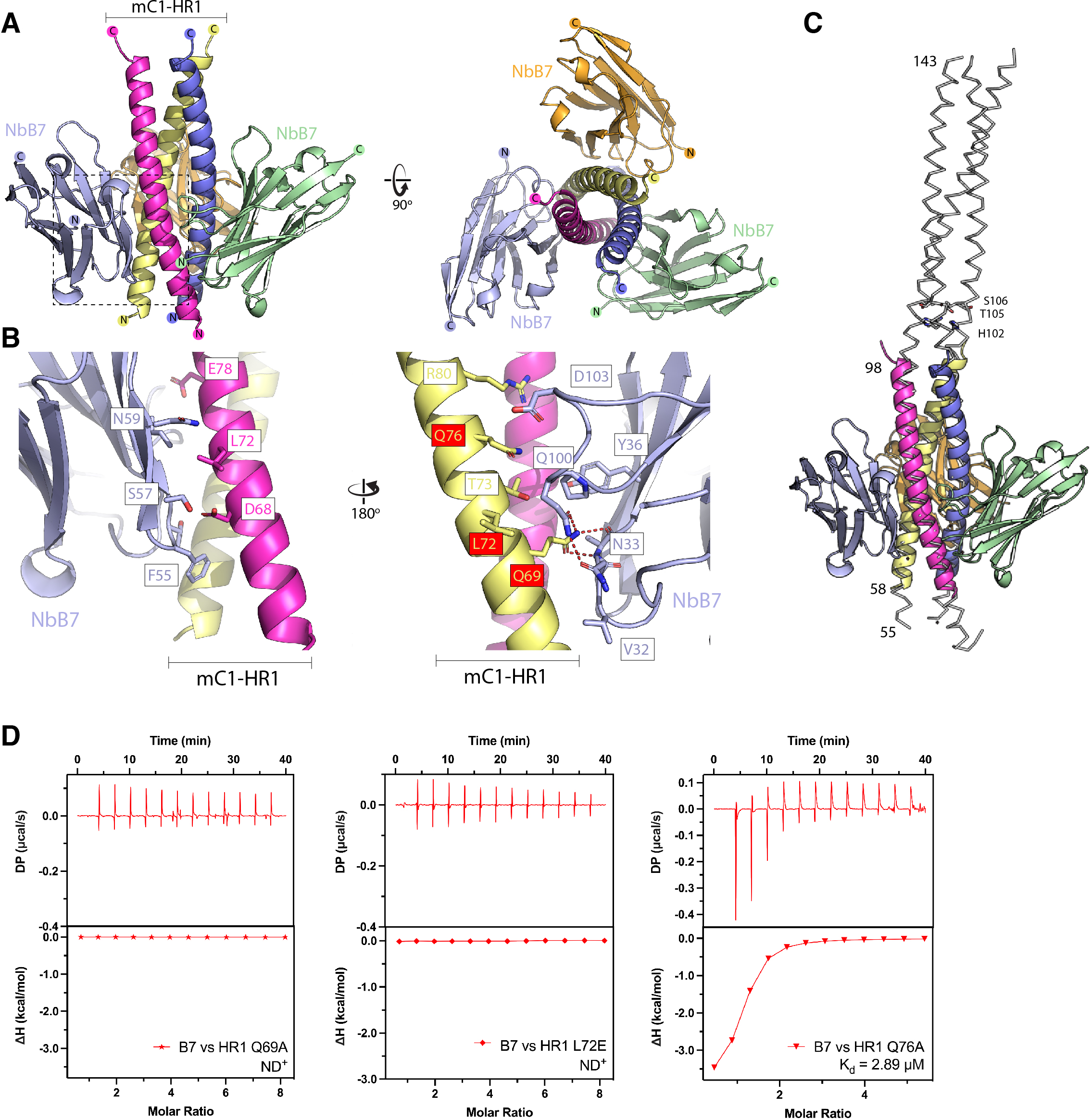
Molecular basis of nanobody NbB7 interaction with mouse Cavin1 HR1 domain. **(A)** The crystal structure of trimeric mC1-HR1 in complex with NbB7. The asymmetric unit contains one copy of the mC1-HR1 protein and one copy of NbB7, with the trimeric structure inferred from the crystallographic symmetry. **(B)** Close-up of the boxed region in (**A**) highlighting key residues mediating interaction between NbB7 (blue) and two adjacent helices of the mC1-HR1 coiled-coil (magenta and yellow). Residues in mC1-HR1 tested by mutagenesis are boxed in red. (**C**) Comparison of the mC1-HR1–NbB7 complex (coloured cartoon) with the previous crystal structure of the mC1-HR1 domain (PDB ID 4QKV)(Kovtun et al., 2014). Numbers on the left indicate N- and C-terminal residues of mC1-HR1 visible in electron density for the NbB7 complex (residues 55-98) and for mC1-HR1 alone (residues 55-143). Residues H102, T105 and S106 mutated in later experiments are indicated on the right. **(D)** ITC binding experiments of NbB7 and selected mutations in mC1-HR1.

Each copy of NbB7 interacts with two adjacent protomers of the mC1-HR1 coiled-coil trimer (**Fig. 4B**). The interaction is, as expected, predominantly mediated by the three CDR loops of the nanobody, but also involves contacts with residues along the face of the β3, β4 and β5 strands of the nanobody structure. Several hydrophobic contacts help to stabilize the interaction including residue Leu72 of mC1-HR1, and Phe55 in NbB7 (within the CDR2 loop). One key contact involves Gln67 of mC1-HR1 which forms a network of hydrogen bonds with the nanobody including with Asn33 in the CDR1 loop and Gln100 in the CDR3 loop. To validate the binding mechanism, we generated several single site mutations in Cavin1 including Q69A, L72E and Q76A and tested their impact on binding by ITC (**Figure 4D; Table S3**). While the Q76A mutation on HR1 significantly reduced the affinity for nanobody NbB7 to micromolar binding affinity both Q69A and L72E reduced the binding to undetectable levels.

To confirm that the structurally mapped sites are responsible for the colocalization with Cavin1 in cells, re-expression of mCherry-Cavin1 in the CRISPR-generated Cavin1 knockout HeLa cells rescued the co-localization of nanobody NbB7 to the plasma membrane (**Figure 5A**). As seen in Cavin1 knockout HeLa cells, Cavin1-Q69A and Q76A mutant still co-localized with NbB7-GFP at the plasma membrane forming the typical puncta (**Figure 5B & 5C**). In contrast, the Cavin1-L72E mutant, which does not bind Cavin1 *in vitro* did not recruit NbB7-GFP for co-localization (**Figure 5D**). These findings further confirm that the leucine residue Leu72 of Cavin1 protein is crucial for the strong binding with nanobody NbB7.

**Figure 5.**
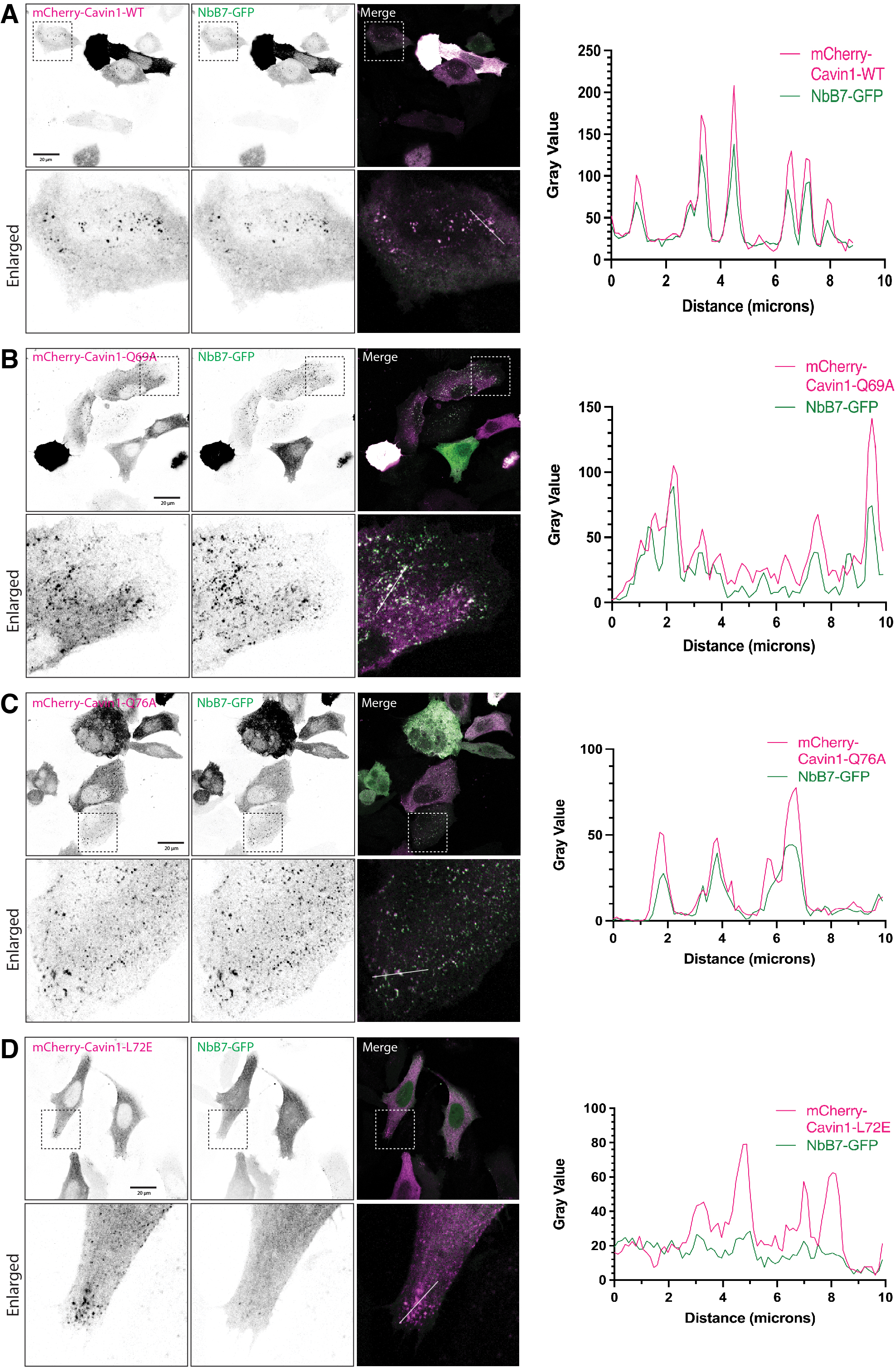
Mutational analysis of the NbB7-Cavin1 interaction in cells. (A-D) Confocal images showing the localization of transiently co-expressed NbB7-GFP nanobody with wild type (WT) or mutated mCherry-Cavin1 (mC-C1) in HeLa Cavin1 knockout cells. Images for each channel were inverted into black and white. Bottom panels display the enlarged images of the selected regions selected in the top panels. Scale bars = 20 μM. Colocalization between NbB7-GFP and mCherry-Cavin1 shown in (**A-D**) was examined by line profile analysis. Scanned lines are indicated in white in the merged images above the curves.

### Structural assessment of the role of a conserved hydrogen-bonded structure in Cavin1 HR1

The observation that the C-terminal half of the Cavin1 HR1 domain is apparently unfolded in the NbB7 co-crystal structure prompted us to investigate this further. There are two previous findings directly relevant to this; firstly, the HR1 domain can be divided into two halves based on a central hydrogen-bonded structure of the coiled-coil (Kovtun et al., 2014), and second there is evidence for unravelling of the C-terminal portion of the HR1 domain upon membrane interaction (Liu et al., 2022). The HR1 domain forms a trimeric coiled-coil of ∼100 residues in length, and is composed of typical heptad repeats, denoted *abcdefg*, where the *a* and *d* positions are generally hydrophobic amino acids such as Leu, Ile and Val required for packing of adjacent helices (Kovtun et al., 2015; Kovtun et al., 2014; Tillu et al., 2018) (**Figure 6A**). However, at the centre of the HR1 domain in Cavin1 are a conserved pair of His and Thr residues (His102 and Thr105 in mouse Cavin1; **Figure 6B**) that form a buried hydrogen-bond interaction. We speculated it could be analogous to the central Arg-Gln pairing in the coiled-coil structures of the soluble SNARE complex (Scales et al., 2001; Sutton et al., 1998). That is, the His-Thr pair in Cavin1 helps to define the specificity of the homomeric and heteromeric protein-protein interactions with itself and Cavin2 and Cavin3 because it prevents binding of other heptad repeat-containing coiled-coil proteins, and may provide a point of instability that allows for regulated assembly and disassembly of the coiled-coil (Kovtun et al., 2015). In addition, Thr104, Thr105 and Ser106 in mouse and human Cavin1 are reported phosphorylation sites in Phosphosite (Hornbeck et al., 2015), and the insulin-stimulated phosphorylation of Thr104, and Ser106 has been reported (Humphrey et al., 2013). The Thr105-Ser106 pair was previously hypothesized to be able to disrupt the hydrogen-bonded pair buried within the coiled-coil (Kovtun et al., 2015).

**Figure 6.**
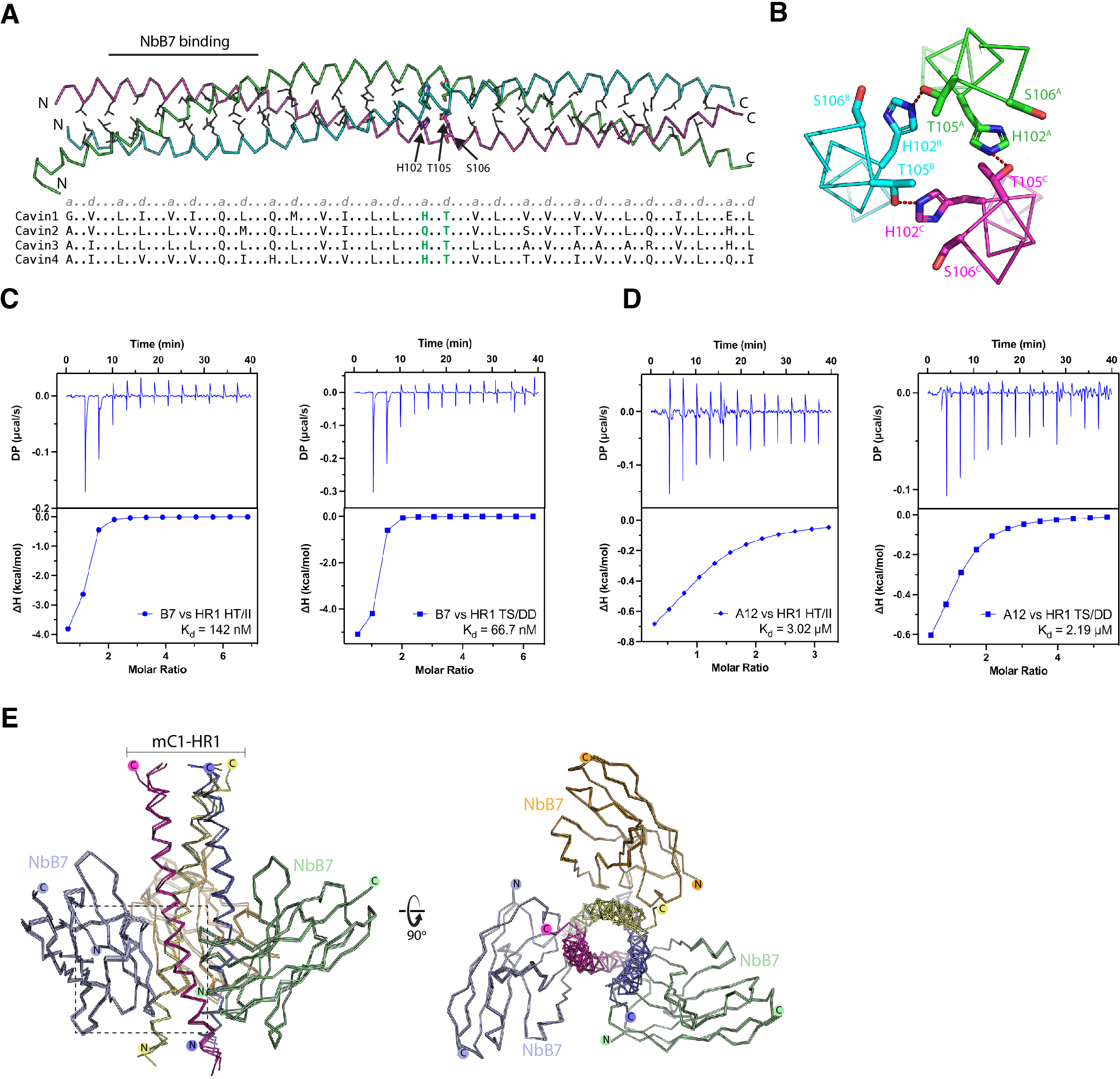
Altered dynamics in binding affinity between nanobodies and hydrogen-bonded pair mutants in Cavin1. **(A)** Heptad repeat sequence alignments of cavin proteins. The top panel represents mouse Cavin1 HR1 structure and the central residues shown with side chains. The bottom panel shows the sequence alignment of heptad repeat in human Cavin1-4 with corresponding residues at position *a* and *d*. The His-Thr (His102-Thr105 in mouse Cavin1) pair was highlighted in dashed box. **(B)** The hydrogen-bonded pair forms between the central His102-Thr105 side chains in mouse Cavin1. **(C)** The interaction of nanobody NbB7 with Cavin1 HR1 HT/II and TS/DD mutants validated by ITC. **(D)** The interaction of nanobody NbA12 with Cavin1 HT1 HT/II and TS/DD mutants validated by ITC. The upper panel represents raw data, and the lower panel represents the binding isotherms. The binding affinity (K_d_) is determined by calculating the mean of at least two independent experiments. **(E)** Structural alignment of the complex NbB7 and Cavin1 HR1 wild-type protein, with HT/II and TS/DD mutants.

To investigate the importance of the His102-Thr105 pair and Thr105-Ser106 we designed two sets of mutations. We hypothesised that altering the His102-Thr105 pair in mouse Cavin1 HR1 domain to Ile sidechains (HT/II mutant) may stabilize this region of the coiled-coil by converting these to typical hydrophobic sidechains of a coiled-coil heptad repeat. The second mutation was to make putative phosphomimetic alterations in the Thr-Ser pair (TS/DD mutant) that we predicted could disrupt coiled-coil formation. Surprisingly, both mutant proteins could be purified similarly to the wild-type mC1-HR1 trimer, and they did not display dramatic differences in stability when we examined their melting temperatures by differential scanning fluorimetry (DSF) (**Figure S2**). When we tested their interactions with NbB7 we found only small changes in their binding affinity and specificity (**Figure 6C-D; Table S4**). The binding affinity (*K*_d_) of NbB7 for the mC1-HR1(HT/II) mutant was moderately enhanced from 520 nM to 142 nM, while its interaction with the mC1-HR1(TS/DD) mutant with nanobody NbB7 was improved by ∼8-fold with a *K_d_* of 66.7 nM. In contrast to NbB7, while NbA12 could still bind the mC1-HR1(HT/II) and mC1-HR1(TS/DD) mutants, the affinities were significantly lower at 3 μM and 2 µM respectively. This suggests that while NbA12 binds an epitope that overlaps with NbB7 (∼residues 65-85) (**Figure 1F**), its binding extends further towards the C-terminus and encompasses the central His102-Thr105 region.

We determined the crystal structures of both mC1-HR1(HT/II) and mC1-HR1(TS/DD) mutants in complex with NbB7, hoping to observe changes in the folding of the C-terminal region of the HR1 domain (**Figure 6E; Table S2**). Interestingly, however, while both structures were obtained in different space groups to the wild-type protein they still showed a lack of density for the C-terminal region of the mC1-HR1 domain beyond residue Lys97. In both cases, the C-terminal region including the mutation sites is therefore still disordered. This also suggests neither the HT/II nor TS/DD mutations have prevented uncoiling of the C-terminal region when bound to the nanobody NbB7 at the N-terminal region.

### Investigating the impact of polar sidechains on cavin-cavin interactions in cells

Although the HT/II and TS/DD mutations had only limited impacts on mC1-HR1 homotrimer stability and folding *in vitro* we discovered a striking phenotype when these mutations are introduced into full-length Cavin1 in cells. To assess if the mutations impacted the ability of Cavin1 to assemble with caveolae or interact with other cavin family members we co-transfected mouse Cavin1-GFP mutants with either Cavin2-mCherry or Cavin3-mCherry in PC3 cells (**Figure 7**). PC3 cells lack expression of any cavin family proteins and do not show morphological caveolae. Caveola formation can be rescued however by exogenous expression of Cavin1, and can recruit both Cavin2 and Cavin3 if they are also co-expressed (Bastiani et al., 2009; Hill et al., 2008). Similar to the wild-type protein (Bastiani et al., 2009; Hill et al., 2008), confocal microscopy showed that Cavin1(HT/II)-GFP and Cavin1(TS/DD)-GFP both localise to typical punctate caveola structures at the plasma membrane, indicating the mutations do not prevent Cavin1’s ability to form caveolae (with membrane embedded CAV1). Intriguingly, however, while Cavin1(HT/II)-GFP can stably recruit both Cavin2-mCherry and Cavin3-mCherry to caveola puncta when co-expressed (**Figure 7A & 7B**), the phosphomimic TS/DD mutant of Cavin1 only recruits Cavin3-mCherry (**Figure 7D**) but does not recruit Cavin2-mCherry which remains dispersed in the cytosol (**Figure 7C**). This striking observation indicates that the putative phosphorylation site in Cavin1 (Humphrey et al., 2013) can play a role in determining specific caveola recruitment of the Cavin2 and Cavin3 homologues.

**Figure 7.**
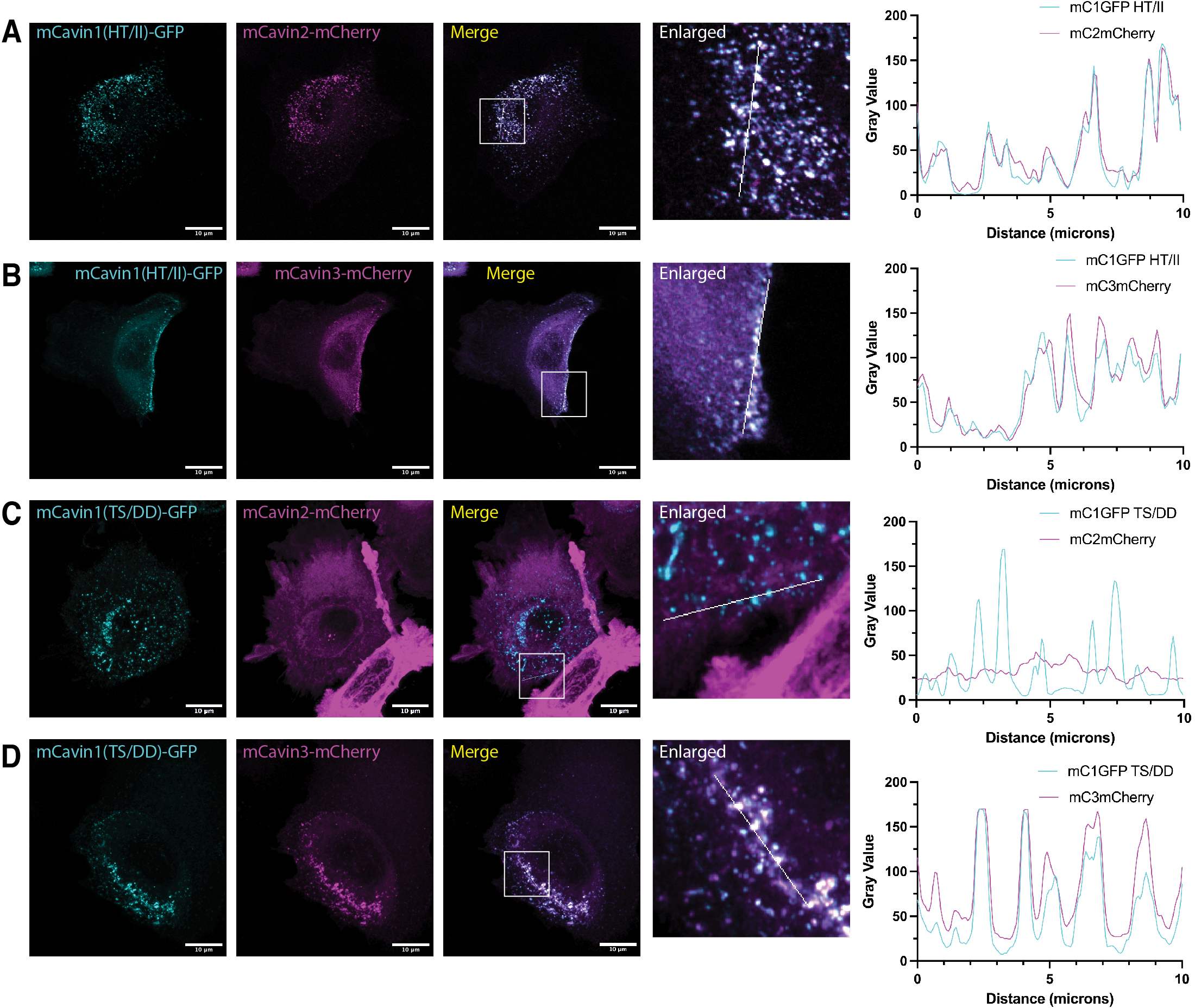
Mutations in the Cavin1 HR1 domain inhibit recruitment of Cavin3 to caveolae. (A-D) Confocal images showing the localization of transiently co-expressed mouse Cavin1-GFP mutants with mouse Cavin2-mCherry or mouse Cavin3-mCherry in PC3 cells. These cells lack endogenous cavins and only form caveolae when Cavin1 is exogenously expressed. Scale bars = 10 μM. Colocalization between Cavin1-GFP and mCherry-tagged proteins was examined by line profile analysis. Scanned lines are indicated in white in the enlarged images.

## Discussion

Cavin1 is an essential component of the caveola coat, interacting with the membrane embedded CAV1 protein, and required for the recruitment of Cavin2 and Cavin3 homologues to the caveola assembly (Bastiani et al., 2009; Gambin et al., 2013; Ludwig et al., 2013; McMahon et al., 2019; Mohan et al., 2015). Its sequence and structure is highly distinctive, with only the HR1 domain known to form an ordered structure in the shape of a membrane-binding trimeric coiled-coil, while the rest of its sequence is either intrinsically disordered or only transiently folded when forming larger complexes (Kovtun et al., 2015; Kovtun et al., 2014; Liu et al., 2022; Tillu et al., 2018; Tillu et al., 2021). Furthermore, molecular dynamics simulations have suggested that the HR1 domain itself may also possess structural plasticity, where its C-terminus can partially unfold to allow for hydrophobic residues to insert into the lipid bilayer (Liu et al., 2022; Lundmark et al., 2024). Although a detailed understanding of the organisation of the core Cavin1 and CAV1 proteins in the caveola coat remains elusive, the structures of the Cavin1 HR1 domain (Kovtun et al., 2014) and of homooligomeric CAV1 discs (Han et al., 2020), and studies of their interactions point towards a heirarchy of multivalent interactions between proteins and the lipid membrane driving caveola formation (Han et al., 2024; Kenworthy et al., 2023; Kovtun et al., 2015; Liu et al., 2022; Lundmark et al., 2024; Mohan et al., 2015; Tillu et al., 2018; Tillu et al., 2021).

To develop new tools for functional analysis and structural studies of Cavin1 we have characterised two nanobodies that bind specifically to the Cavin1 HR1 coiled-coil. Nanobodies and their derivatives are rapidly emerging as powerful research tools for structural determination, cellular engineering in biological research and as potential biologic therapeutics (Muyldermans, 2021; Uchański et al., 2020). The two nanobodies each bind with similar affinities to the trimeric HR1 domain, with overlapping but not identical epitopes within the N-terminal half of the extended coiled-coil structure. They are relatively specific and bind to Cavin1 from zebrafish, although they show some mild cross-reactivity with Cavin2 and Cavin4. In the presence of both nanobodies *in vitro*, Cavin1 was found to retain its inherent anionic membrane interaction and membrane remodelling ability. In the future cryo-electron tomography of the membrane tubules generated by the Cavin1-Nb complexes may reveal the precise mode of binding of the membrane by the Cavin1 protein and whether the presence of the nanobodies affects the conformation of Cavin1 upon membrane recognition.

The crystal structure of NbB7 in complex with mC1-HR1 revealed the precise mechanism of its interaction and allowed us to confirm the binding site via structure-based mutagenesis. However, a remarkable feature of this crystal structure was that the C-terminal portion of the coiled-coil of HR1 (residues 98-155) was unexpectedly absent in the electron density compared to mC1-HR1 on its own (Kovtun et al., 2014). Based on considerations of the symmetry of the crystal lattice, and the fact that structures of two other mC1-HR1 mutants in complex with NbB7 show a similar lack of electron density, we are confident that this indicates the C-terminal residues have become disordered due to unravelling of the coiled-coil in this region. What does this mean? We believe this points to an inherent instability in the HR1 coiled-coil beyond the central hydrogen-bonded His and Thr residues, and that either upon binding of NbB7 at the N-terminus, or during crystallisation of the complex, the C-terminal region can transition into an unfolded state or shifts its equilibrium between folding and unfolding. Although this remains to be categorically proven, the observation provides support for the previous modelling that indicates the Cavin1 HR1 domain can partially unfold at its C-terminus when binding to a lipid membrane (Liu et al., 2022).

The discovery that the C-terminal half of the Cavin1 HR1 domain can adopt an unfolded state, made possible using NbB7, prompted us to examine the site of ‘uncoiling’ more closely. The region corresponds to the centre of the coiled-coil defined by the hydrogen-bonded residues His102 and Thr105 (Liu et al., 2022), and a site of putative phosphorylation including Thr105 and Ser106 that are upregulated in insulin-stimulated adipocytes (Humphrey et al., 2013). We hypothesised that mutations at these sites might act to either stabilise the HR1 coiled-coil (substituting the buried hydrogen-bond pair with hydrophobic side chains) or perturb its assembly (adding negative charges to the Thr and Ser side-chains). These mutations did not dramatically impact the coiled-coil stability *in vitro* nor the ability of the full-length Cavin1 protein to promote caveola formation in cells. However, without affecting Cavin3, the phosphomimetic mutation in Cavin1 led to a complete inability to recruit the Cavin2 homologue to caveolae. Previous studies have shown that Cavin1 forms distinct complexes with either Cavin2 or Cavin3 (Bastiani et al., 2009; Gambin et al., 2013; Ludwig et al., 2013; McMahon et al., 2019; Mohan et al., 2015), and that Cavin3 can be specifically released from caveolae upon cellular stress (McMahon et al., 2021). Our results show that sequences within the central region of the HR1 domain can define specificity for Cavin3 over Cavin2, and potentially, that post-translational modification of Cavin1 at this site could be important for regulating these specific interactions in the cell. This raises the fascinating possibility that the cavin complement, and presumably function, of caveolae can be dynamically modulated by post-translational modifications *in vivo* in response to specific physiological signals.

This work provides the first report of structurally characterised nanobody probes that can be used as molecular tools for studying caveola associated proteins. They have provided novel insights into the folding and stability of the Cavin1 HR1 domain, and NbB7 can be used for the imaging of Cavin1 and caveolae in cells by fluorescence and electron microscopy. and imaging Cavin1 *in situ* and *in vivo.* For example, in other work we have recently applied NbB7 to quantify changes in caveolae numbers under different lipid peroxidation conditions (Wu et al., 2024). Future applications could include use as affinity reagents for proteomics of endogenous Cavin1, altering Cavin1 activity via over-expression or targeting of modifying enzymes, use as fiducial markers in cryoelectron tomography studies of Cavin1-coated membranes and vesicles, for studying the dynamics of caveolae by high-resolution caveola tracking in cells, or for correlative light and EM (CLEM) imaging of caveolae to answer many long-standing mysteries about their organization.

## Materials and Method

### Molecular cloning and plasmids

All plasmids are listed in **Table S5**. For bacterial expression of cavin proteins and their helical domains, DNA encoding N-terminal 6x Histidine-Ubiquitin (His-Ub) tagged mouse Cavin1 HR1, shorter fragment of mC1 HR1 (residues 45-103), mC1 HR1 5Q mutant, mouse Cavin2 (mC2) HR1, zebrafish Cavin1a (zC1a) HR1, zebrafish Cavin4a (zC4a) HR1, zebrafish Cavin4b (zC4b) HR1, single-site mutants of mC1 HR1 including mC1 HR1 Q69A, L72E, Q76A were all cloned into pHUE vector at *SacII* restriction enzyme site using overlap extension-polymerase chain reaction (OE-PCR) (Catanzariti et al., 2004). DNA encoding mouse Cavin1 full-length was cloned into 2K-T vector containing a His-MBP tag at *SspI* restriction enzyme site using OE-PCR. The N-terminal GST-tagged mouse Cavin1 HR1, and I102-I105 (HT/II), D105-D106 (TS/DD) construct was cloned into pGEX-4T2 vector with site-specific thrombin cleavage sequence using OE-PCR. For mammalian cell expression, nanobody NbA12 and NbB7, mouse Cavin1 full-length point mutant Q69A, L72E, Q76A, I102-I105 (HT/II), D105-D106 (TS/DD) were cloned into pEGFP-N1 vector with a C-terminal GFP tag at the *BamHI* restriction enzyme site. Site-directed mutagenesis was applied to all mutant constructs with custom designed primers.

### Recombinant protein expression and purification of cavin proteins

N-terminal His-Ub tagged mC1 HR1, shorter fragment of mC1 HR1 (residues 45-103), mC1 HR1 5Q mutant, mC2 HR1, single-site mutants of mC1 HR1 including mC1 HR1 Q69A, mC1 HR1 L72E, mC1 HR1 Q76A, and His-MBP tagged full-length mC1 were transformed into *E. coli* BL21(DE3)pLysS competent cells by heat shock at 42°C. The codon-optimized zC1a HR1, zC4a HR1, zC4b HR1 and N-terminal GST-tagged mC1 HR1, I102-I105 (HT/II), D105-D106 (TS/DD) mutant were transformed into *E. coli* BL21 CodonPlus™ (DE3) cell. Cells were propagated in fresh LB broth at 37°C at 200 rpm until OD_600_ reaching between 0.8 - 1.0, and induced with 0.5 mM Isopropyl ß-D-1-thiogalactopyranoside (IPTG) at 18°C for 16 h. Cells were harvested and lysed using a continuous flow cell disruptor (Constant Systems Limited, UK) at 25-32 kPsi pressure in 20 mM HEPES pH 7.4, 500 mM NaCl buffer (GF500) with 50 μg/ml benzamidine hydrochloride, 10 μg/ml deoxyribonuclease I (DNase I), 0.5% w/v Triton X-100 and addition of 5 mM Imidazole (His-UB tagged cavins only) followed by high-speed centrifugation 38,000 × g for 30 min. All recombinant proteins with a His-UB tag and His-MBP full-length mC1 were purified by TALON^®^ metal affinity resin (ClonTech, Scientifix Cat No. 635503) and washed with buffer GF500 containing 5 mM Imidazole to remove contaminants (used for ITC experiments). Protein samples were then eluted by GF500 containing 300 mM Imidazole. GST-tagged mouse Cavin1 HR1 domain, and I102-I105 (HT/II), D105-D106 (TS/DD) mutant were purified by glutathione Sepharose^™^ affinity resin (GE healthcare) and GST-tag was removed on column by 100 units/ml thrombin (Sigma-Aldrich) at 4°C and eluted by GF500 buffer. The eluted proteins were subsequently subjected to HiLoad^™^ 16/600 Superdex^™^ 200 prep grade column (GE Healthcare) using a semi-automated ÄKTA Purifier FPLC purification system (GE Healthcare) in filtered 20 mM HEPES pH 7.4, 150 mM NaCl buffer (GF150) (for ITC experiments) or GF500 buffer in addition of 2 mM dithiothreitol (DTT; for crystallization).

### Generation of nanobodies

An alpaca was immunized six times with 200 ug of purified recombinant mouse Cavin1 HR1. The adjuvant used was GERBU FAMA. Immunization and handling of the alpacas for scientific purposes was approved by Agriculture Victoria, Wildlife & Small Institutions Animal Ethics Committee, project approval No. 26-17. Blood was collected three days after the last immunization for the preparation of lymphocytes. Nanobody library construction was carried out according to established methods as described (Pardon et al., Nature Protocols 2014). Briefly, alpaca lymphocyte mRNA was extracted and amplified by RT-PCR with specific primers to generate a cDNA nanobody library. The library was cloned into a pMES4 phagemid vector, amplified in *E. coli* TG1 strain and subsequently infected with M13K07 helper phage for recombinant phage expression.

Biopanning for recombinant mouse Cavin1 HR1 nanobodies using phage display was performed as previously described with following modifications (Pardon et al, Nature Protocols 2014). Phages displaying mouse Cavin1 HR1 -specific nanobodies were enriched after two rounds of biopanning on immobilised mouse Cavin1 HR1. After the second round of panning, we screened single clones and positive clones were selected for further analysis. 43 positive clones were sequenced and 7 distinct nanobody clonal groups were identified based on unique CDR3 sequences.

### Recombinant protein expression and purification of nanobody

Nanobodies were expressed in *Escherichia coli* WK6 cells and purified as previously described (Pardon et al., Nature Protocols 2014). Nanobodies were cloned with a C-terminal 6x Histidine tag into pMES4 expression vector for periplasmic expression in *E. coli* WK6 (*Su*^−^) (Zell and Fritz, 1987). Nanobodies were expressed in TB media supplemented with 100 mg/ml ampicillin, 20% glucose and 2mM MgCl_2_ and then induced with 1 mM IPTG at 25°C for 20 h. To extract proteins from the periplasm by osmotic shock, the harvested cell pellets were resuspended in 200 ml TES buffer containing 0.2 M Tris-HCl pH 8.0, 0.5 mM EDTA, 0.5 M sucrose with addition of 50 mg/ml benzamidine hydrochloride and incubated on an orbital shaking platform at 4°C overnight. 400 ml of 1/4 TES buffer was then added to the resuspended pellet and shaken for another 1 h at 4°C. The lysates were centrifuged in high speed at 14,000 ×g for 30 min to extract proteins. His-tagged nanobodies were purified by TALON^®^ metal affinity resin and then subjected to size exclusion chromatography in a HiLoad^™^ 16/600 Superdex^™^ 200 prep grade column (GE Healthcare) using a semi-automated ÄKTA Purifier FPLC purification system (GE Healthcare).

### GST-pulldown assay

GST-pulldown assay was carried out using GST-tagged mouse Cavin 1 HR1 domain as bait protein for His-tagged Cavin1-specific nanobodies, A4, NbA12, NbB7, D5, D6, H3 and H5. 1 nmol of GST-tagged mouse Cavin1 HR1 and GST alone were mixed with 2 nmol of His-tagged Cavin 1-selected nanobodies in 500 *μ*l of pulldown buffer (20 mM HEPES, 150 mM NaCl, 1 mM DTT, 0.1% IGEPAL, pH 7.4), respectively. The protein mixture was incubated in the arbitrary shaking platform for 1 h at 4 °C. Then the mixture was centrifuged at 17,000 rpm for 20 min to remove any precipitated proteins. The clarified protein mixture and GST alone were incubated with 50 μl of pre-equilibrated glutathione Sepharose^™^ affinity resin (GE healthcare), respectively. The reaction mixtures were incubated at 4 °C for 1 h and the protein bound glutathione Sepharose^™^ affinity resin (GE healthcare) was spun down at 5,000 rpm for 30 s. After removing the supernatant, and the resin was washed 4 times with pulldown buffer. Resin was then mixed with 50 μl of SDS sample buffer and analysed by SDS-PAGE.

### Isothermal titration calorimetry

All microcalorimetry experiments were conducted in GF150 buffer at 25°C using a MicroCal PEAQ-ITC (Malvern). ITC experiments were performed with one single 0.4 μl injection followed by 12 injections of 3.22 μl each with 180 s injection spacing. Nanobody NbA12 at 500 μM and NbB7 at 400 μM were titrated into 30 μM monomeric HR1 domain of cavins and His-MBP full-length mC1, respectively. The interaction of mouse Cavin1 HR1 domain and nanobody NbB7 in the presence of competing nanobody NbA12 was conducted by titrating 1 mM nanobody NbB7 into 50 μM mC1 HR1 pre-incubated with 50 μM nanobody NbA12. 200 μM nanobody NbA12 or NbB7 were titrated into 6.25 μM trimeric mouse Cavin1 HR1 I102-I105 (HT/II), D105-D106 (TS/DD) mutant. Data was analyzed by Malvern software by fitting and normalized to a single-site binding stoichiometry and thermodynamic profiles were presented by Prism 8.0.1 version (GraphPad Software). Experiments were conducted with at least three technical replicates for data reproducibility.

### Crystallisation and data collection

Hanging-drop vapor diffusion method was applied for crystallization screening under a 96-well plate by Mosquito Liquid Handling robot with a 1:1 ratio of protein and reservoir solution at 20°C. To co-crystallize nanobody with mouse Cavin1 HR1 domain, 1.5-fold molar excess of purified nanobody NbA12 and NbB7 were incubated separately to the purified mouse Cavin1 HR1 domain and undergo subsequent size exclusion chromatography to form a complex with a final concentration of 12.6 mg/ml. Initial crystals of nanobody NbB7-mouse Cavin1 HR1 complex were obtained in many commercial screen conditions but the best diffraction quality crystal was optimized in 0.65 M potassium thiocyanate, 19.5% PEG 2000 MME.

For co-crystallization of mouse Cavin1 HR1 I102-I105 (HT/II), D105-D106 (TS/DD) mutants with nanobody NbB7, the GST-tag on HR1 mutants was cleaved by thrombin and pre-incubated with 1.5-fold molar excess affinity-purified nanobody NbB7 overnight to form a complex. The protein complex crystals were produced using the hanging drop method in the crystallization condition containing 0.2 M potassium thiocyanate, 20% PEG 3350 and 0.2 M potassium citrate, 20% PEG 3350. All the crystals were soaked in the appropriate cryoprotectant solutions containing the crystallization solutions supplemented with 25% glycerol and flash-cooled in a nitrogen gas stream for transporting to the Australian Synchrotron for data collection. X-ray diffraction data was measured on the MX2 beamline at the Australian Synchrotron.

### Crystal structure determination

The collected data was integrated using XDS (Kabsch, 2010) and scaled by AIMLESS (Evans and Murshudov, 2013). The crystal structure of nanobody NbB7 in complex with mouse Cavin1 HR1, and I102-I105 (HT/II), D105-D106 (TS/DD) mutants were solved by molecular replacement using PHASER (McCoy et al., 2007) with the available mouse Cavin1 HR1 domain [Protein Data Bank (PDB) ID: 4QKV] and GFP-nanobody [PDB ID: 3OGO] as the initial model. The initial electron density map was then refined in PHENIX suite (Adams et al., 2010) and rebuilt in COOT. Data and refinement statistics are summarized in **Table S2** and molecular figures are presented using PyMOL.

### Liposome preparation and pelleting assay

Folch liposomes were freshly prepared by using 100% Folch fraction I (Sigma-Aldrich) in chloroform to generate a lipid film in a mini-round bottom flask and dried gently under a nitrogen stream and then under vacuum overnight. To form multilamellar vesicles (MLVs), the dried lipid film was rehydrated in GF150 buffer followed by 10 rapid freeze-thaw cycles using acetone in dry ice and 60°C water. The pelleting assay was conducted in a total volume of 100 μl comprising 50 μl of 1mg/ml MLVs and 50 μl of 10 μM GST-mC1 HR1, 10 μM nanobody NbA12 or NbB7, 10 μM GST-mC1 HR1 in complex with nanobody NbA12 or NbB7 in a 1-to-1 ratio. The reaction mixtures were incubated at 25°C for 10 min followed by centrifugation at 60,000 × g for 15 min at 4°C in Optima^™^ MAX-XP tabletop ultracentrifuge (Beckman Coulter) using TLA100 rotor. The supernatant was collected, and the pellet was then carefully separated from supernatant resuspending in 50 μl GF150 buffer containing 4 × loading dye before analyzed by SDS-PAGE.

### Liposome preparation and tubulation assay

Liposome tubulation assay was perfumed as described previously (Kovtun et al., 2014). Liposomes were prepared as described above and rehydrated in GF150 buffer followed by 3 repetitive freeze-thaw cycles to form multilamellar layers. Large unilamellar lipid vesicles (LUVs) were generated by subjecting the liposomes to 21 rounds of extrusion through an 800-nm polycarbonate membrane using an Avanti miniextruder. Purified complexes of nanobody NbA12 or NbB7 pre-incubated with His-MBP full-length mouse Cavin1 at 0.1 mg/ml were incubated with 0.2 mg/ml liposomes for 2 min at 25°C, respectively. Samples were then immediately applied onto formvar carbon-coated electron microscopy grids (300 mesh) for 10 s. Excess samples were eliminated by blotting with Whatman filter paper at corner. The grids were washed with distilled water for 3 times before application of 2% uranyl acetate stain. The grids were blotted again to remove excess stain and left to air dry before viewing. Images were captured using a JEOL 1011 transmission electron microscope at 80 kV.

### Cell culture and transfection

Cell lines were sourced from the American Type Culture Collection (ATCC). HeLa cells, A431 cells and baby hamster kidney (BHK) cells were maintained in Dulbecco’s modified eagle medium (DMEM; Thermo Fisher Scientific) supplemented with 10% fetal bovine serum and 2 mM L-glutamine (Thermo Fisher Scientific) at 37°C. On the following day, HeLa cells, A431 cells, and BHK cells were transfected with Lipofectamine 2000 (Themo Fisher Scientific) at 70% confluency. Cavin1 knockout HeLa cells were transfected with Lipofectamine 3000 (Thermo Fisher Scientific) at 70% confluency according to the manufacturer’s instructions.

### Co-immunoprecipitation

NHS-activated Sepharose^™^ 4 fast flow (Cytiva) resin was prepared by washing with GF500 buffer and coupled with GFP-nanobody (NbA12/NbB7) with overnight rotation using the manufacturer’s protocol. The unbound GFP-nanobody was removed by washing with GF500 buffer and the resin blocked by 0.2 M Ethanol amide for 5 h. The coupled resin was washed 2 times with GF500 buffer before storage in PBS at 4 °C.

HeLa cells were cultured on Corning^®^ tissue-culture dishes until reaching a 50-70% confluency. Cells were removed by mechanical scraping from dishes and harvested by centrifugation at 500 × g for 10 min. The harvested cells were lysed in ice-cold GF150 buffer with 0.5% Triton X-100, 0.1 mg/mL Benzamidine, 10 μg/mL DNase I, phosphatase inhibitor cocktail (Roche PhosSTOP^™^) and protease inhibitor cocktail (COMPLETE^™^ EDTA-free). They were lysed by sonication using ultrasonic cell disruptor with 3 pulses at a time on ice and centrifuged at 1,000 × g for 2 min. Supernatant was collected from cell lysates and 100 μL of supernatant was boiled with 40 μL 4 × SDS-PAGE buffer as lysate control for analysis. The rest of the supernatant was subjected to nanobody NbA12/NbB7 coupled NHS-activated Sepharose^™^ resin for 2 h incubation at 4 °C. The resultant pellet was then washed with ice-cold GF150 buffer 3 times and then suspended in 200 μL of GF150 buffer with 60 μL 4 × SDS-PAGE buffer for Western blot analysis.

Whole-cell lysates were collected from HeLa cells grown in the 6-well dishes and lysed by RIPA buffer (150 mM NaCl, 0.5% Triton X-100, 0.1% SDS, 50 mM Tris pH 7.4), supplemented with a phosphatase inhibitor cocktail (Roche PhosSTOP^™^) and protease inhibitor cocktail (COMPLETE^™^ EDTA-free) at 4 °C. After scratching from the bottom of 6-well plate and vortex to homogenise, the cell lysates were incubated on ice for 5 min and undergone centrifugation at 10,000 × g for 3 min at 4 °C. The supernatant was collected and boiled with 4 × SDS-PAGE buffer at 95 °C for 3 min for sample collection.

### Western blotting

Samples were separated on SDS-PAGE gel in 1 × MES buffer at 90 V for the first 20 mins and 130 V until the dye-front reached the bottom of the gel. The proteins were transferred from SDS-PAGE gels to polyvinylidene difluoride membrane (PVDF) via dry transfer according to the Abcam’s Western blot protocol. The PVDF membrane was blocked by incubating with 5 % skim milk in PBS containing 0.1 % Tween-20 (PBST) for 1 h at room temperature to prevent non-specific binding. The PVDF membrane was incubated with anti-GFP (Polyclonal mouse, Proteintech^®^, dilution 1:1000), anti-Cavin1 (Polyclonal rabbit, Proteintech^®^, dilution 1:1000), anti-CAV1 (Polyclonal rabbit, BD Transduction Laboratories, dilution 1:1000) primary antibodies were incubated with the PVDF membrane in 5% skim milk with PBST overnight at 4 °C. The membrane was washed with PBST for 4 x 5 mins and then incubated with appropriate secondary antibodies (dilution 1:2000 for horseradish peroxidase-conjugated goat anti-mouse IgG, 1:12000 for horseradish peroxidase-conjugated goat anti-rabbit IgG) in PBST with 5% skim milk for 90 min. The membrane was washed again with PBST for three times and developed using chemiluminescence-based immunodetection by Novex^®^ chemiluminescent substrate (Invitrogen). Images were then obtained by Hi-resolution blot imaging and normal image scanning for colorimetric detection following the manufacturer’s instructions.

### Immunofluorescence imaging and confocal microscopy

Cells expressing GFP-nanobody NbB7 or NbA12 constructs were plated on glass coverslips and fixed with 4% paraformaldehyde in PBS for 30 min at room temperature. They were then washed with PBS and permeabilized with 0.1% Triton-X 100 in PBS followed by addition of 2% bovine serum albumin (BSA) for 1 h blocking. Cells were probed with primary antibody anti-Cavin1 (Polyclonal rabbit, Proteintech^®^, dilution 1:800) or anti-CAV1 (Polyclonal rabbit, BD Transduction Laboratories, dilution 1:800) for 45 min at 4 °C. Secondary antibodies goat anti-rabbit IgG (dilution 1:400) were then incubated on coverslips for 45 min at room temperature. The coverslips were mounted in Mowiol in PBS. Images were acquired using an Inverted LSM 880 Fast Airyscan Microscope equipped with × 63 oil immersion objective lens.

Cavin1 knockout HeLa cells were seeded onto glass coverslips and fixed with 4% paraformaldehyde in PBS for 15 min at room temperature. Following fixation, cells were washed three times with PBS. Coverslips were then mounted in Mowiol in 0.2 M Tris-HCl, pH 8.5. Images were captured using a ZEISS LSM 880 Confocal Microscope. Intensity line scan analysis was conducted in ImageJ 2.0 using a “line” tool to draw a line at the measurement site in the first channel and repeat it in the second channel to measure fluorescence intensities across the line in both channels using “plot profile” option.

### APEX labelling electron microscopy

BHK cells were seeded into 35 mm tissue culture dishes overnight and then transfected with Lipofectamine 3000 as per manufacturer’s instructions. Dishes were fixed 24 h later with 2.5% glutaraldehyde in PBS and then washed repeatedly in PBS. Fixed cells were treated with a freshly prepared solution of 0.05% DAB in PBS for 10 min followed by incubation in a solution of 0.05% DAB containing 0.01% H_2_O_2_ for 30 min at room temperature. Cells were postfixed with 1% osmium for 2 min and underwent a series of dehydration steps using increasing percentages of ethanol. Cells were then subjected to a series of infiltration steps with LX112 resin in a Pelco Biowave system followed by 24 h incubation at 60°C to polymerize. Ultrathin sections were attained on a ultramicrotome (UC6, Leica) and imaged using a JEOL1011 transmission electron microscope at 80 kV.

### Differential scanning fluorimetry

Thermal unfolding assay was conducted using a ViiA7 real-time PCR instrument (Applied Biosystems) to detect preferential binding of a fluorophore to unfolded molecules. Briefly, 2 mg/ml of freshly purified mouse Cavin1 HR1 (control), HT/II and TS/DD mutant were centrifuged at 17,000 RPM at 4°C for 15 min to eliminate possible precipitants and then mixed with freshly prepared 500X SYPRO orange dye (Life Sciences) to a final concentration of 8 μM for thermal denaturation measurement. The sample mixture was then loaded onto a 96-well plate and experiments were performed in four replicates. Relative fluorescence units were measured from 24°C to 80°C with 1°C increment using the ROX dye calibration. The melting temperature (Tm) was determined by fitting the data into a Boltzmann sigmoidal curve in Prism version 10.2.0 (GraphPad Software).

## ACKNOWLEDGEMENTS

We acknowledge use of the University of Queensland Remote Operation Crystallization and X-ray (UQ ROCX) Facility and the assistance of G.King and K.A.Arachchige. X-ray data were collected on the MX1 and MX2 microfocus beamline at the Australian Synchrotron. The authors acknowledge the use of the Microscopy Australia Research Facility at the Centre for Microscopy and Microanalysis at The University of Queensland.

## Author contributions

Conceptualization, YG, VAT, BMC;

Methodology, YG, VAT, YPW, JR, TEH, KEC, SW, QG, EL, WHT, RGP, BMC;

Investigation, YG, VAT, BMC, JR, YPW, TEH;

Writing – Original Draft, YG, VAT, RGP, BMC;

Writing – Review & Editing, YG, VAT, WHT, RGP, BMC;

Funding Acquisition, WHT, RGP, BMC;

Supervision, VAT, RGP, BMC.

## Data Availability

All data related to this study is available from the corresponding authors on request.

## Supplementary Information

**Figure S1.**
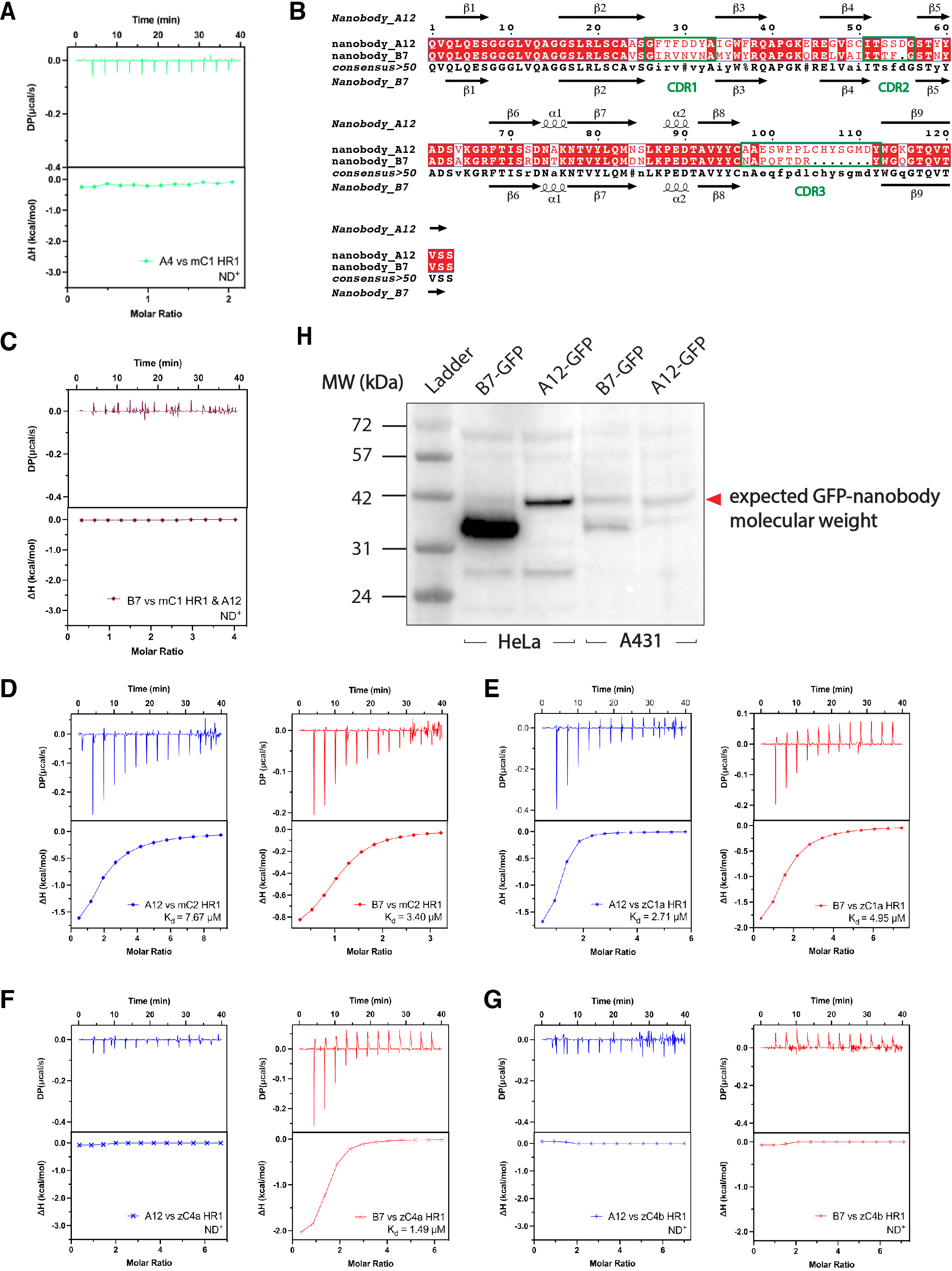
Nanobody sequences, binding *in vitro*, and expression in cells. **(A)** ITC experiment showing lack of affinity between NbA4 and mC1-HR1. (**B**) Sequence alignment of nanobody NbA12 and NbB7. The three CDRs are highlighted in green boxes. **(C)** Nanobody NbA12 and NbB7 competes for binding with mouse Cavin1 HR1. ND^+^: no binding detected. **(D)** Binding of NbA12 and NbB7 with mouse Cavin2 HR1 domain measured by ITC. **(E)** Binding of both nanobodies with zebrafish Cavin1a HR1 domain measured by ITC. **(F)** Binding of nanobody NbB7 with zebrafish Cavin4a HR1 domain was measured by ITC but no binding was observed for nanobody NbA12. **(G)** ITC thermograms showing that both nanobodies could not interact with zebrafish Cavin4b HR1 domain. ND^+^: no binding detected. The upper panel represents raw data, and the lower panel represents the normalized and integrated binding isotherms fitted with a 1-to-1 binding ratio. The binding affinity (K_d_) is determined by calculating the mean of at least two independent experiments. (**H**) Western blot with anti-GFP antibody to detect expression of NbB7-GFP and NbA12-GFP in HeLa cells and A431 cells. Expression levels are higher in HeLa cells, and suggest that some nicking of the NbB7 protein occurs at higher expression levels.

**Figure S2.**
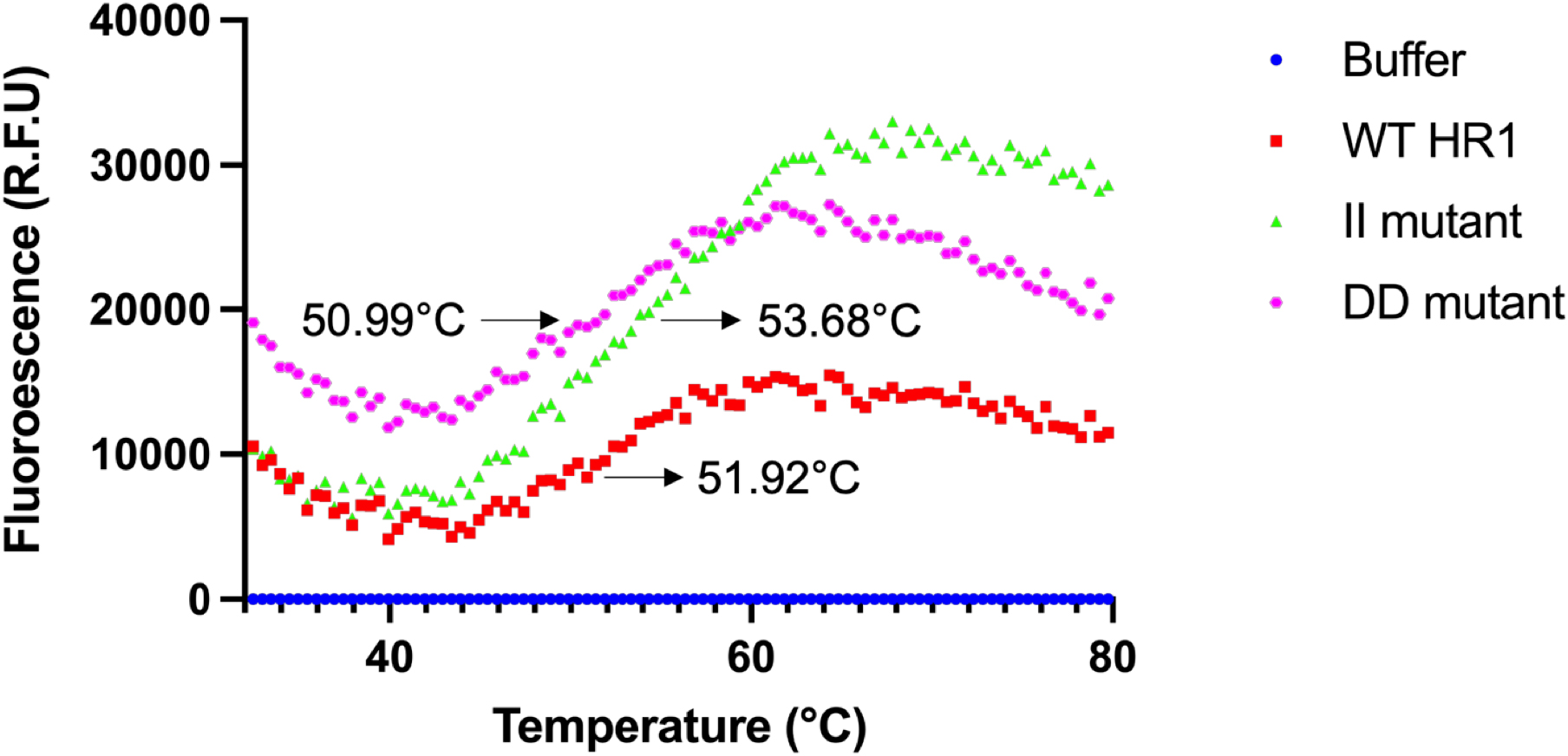
Thermal stability of Cavin1 HR1 domain mutants. **(A)** Differential scanning fluorimetry (DSF) assay to measure the melting temperature of the mC1-HR1 trimer and comparison with the HT/II and TS/DD mutants.

**Table S1.** Thermodynamic parameters for the interaction of cavin proteins with nanobodies by ITC.

|  | K<sub>d</sub> (μM) | ΔH (kcal/mol) | ΔG (kcal/mol) | -TΔS (kcal/mol) |
| --- | --- | --- | --- | --- |
| NanobodyNbA12 |  |  |  |  |
| mC1 HR1 | 0.50 ± 0.14 | -5.70 ± 0.08 | -8.57 ± 0.16 | -2.88 ± 0.23 |
| mC1 HR1 45-103 | 0.57 ± 0.00 | -5.07 ± 0.09 | -8.54 ± 0.01 | -3.35 ± 0.07 |
| mC1 HR1 5Q | 0.14 ± 0.02 | -4.09 ± 0.83 | -9.35 ± 0.11 | -5.26 ± 0.73 |
| mC1 full-length | 8.88 ± 1.73 | -0.48 ± 0.09 | -6.90 ± 0.11 | -6.42 ± 0.21 |
| mC2 HR1 | 7.67 ± 6.55 | -1.74 ± 0.23 | -7.12 ± 0.59 | -5.38 ± 0.82 |
| zC1a HR1 | 2.71 ± 2.38 | -2.75 ± 1.25 | -7.75 ± 0.62 | -5.00 ± 1.86 |
| zC4a HR1 | No binding detected |  |  |  |
| zC4b HR1 | No binding detected |  |  |  |
| Nanobody NbB7 |  |  |  |  |
| mC1 HR1 | 0.52 ± 0.35 | -4.21 ± 0.49 | -8.67 ± 0.45 | -4.45 ± 0.93 |
| mC1 45-103 | 0.45 ± 0.14 | -4.77 ± 0.52 | -8.68 ± 0.19 | -3.91 ± 0.71 |
| mC1 HR1 5Q | 0.89 ± 0.02 | - 4.45 ± 1.09 | -8.26 ± 0.01 | -3.81 ± 1.11 |
| mC1 full-length | 1.73 ± 0.18 | -6.39 ± 0.52 | -7.87 ± 0.06 | -1.48 ± 0.58 |
| mC2 HR1 | 3.40 ± 2.46 | -0.78 ± 0.09 | -7.56 ± 0.47 | -6.77 ± 0.38 |
| zC1a HR1 | 4.95 ± 2.47 | -2.01 ± 1.46 | -7.28 ± 0.31 | -5.28 ± 1.76 |
| zC4a HR1 | 1.64 ± 0.21 | -2.26 ± 0.05 | -7.90 ± 0.07 | -5.65 ± 0.02 |
| zC4b HR1 | No binding detected |  |  |  |
| NbB7 vs mC1 HR1 & NbA12 |  | No binding detected |  |  |

**Table S2.** Statistics summary of X-ray crystallographic structure determination.

|  | Nanobody NbB7-mouse Cavin1 HR1 | Nanobody NbB7-mouse Cavin1 HR1<br>HT/II | Nanobody NbB7-mouse Cavin1 HR1<br>TS/DD |
| --- | --- | --- | --- |
| **Data collection statistics** |  |  |  |
| PDB ID | 9EGN | 9EG6 | 9EIU |
| Space group | P63 | C 1 2 1 | P 3 |
| Resolution (Å) | 46.50 – 1.57 | 48.82 – 3.2 | 30.6-4.0 |
| a, b, c (Å) | 58.90, 58.90, 113.12 | 102.32, 59.24, 133.82 | 58.67, 58.67, 61.29 |
| α, β, γ (°) | 90.0, 90.0, 120.0 | 90.0, 97.48, 90.0 | 90.0, 90.0, 120.0 |
| Total observations | 641590 (27806) | 87884 (15457) | 10543 (3143) |
| Unique reflections | 31084 (1492) | 13349 (2402) | 1975 (572) |
| Completeness (%) | 99.7 (95.0) | 99.8 (100.0) | 99.8 (99.8) |
| R<sub>merge</sub><sup>+</sup> | 0.045 (0.469) | 0.297 (1.281) | 0.076 (0.159) |
| R<sub>pim</sub><sup>*</sup> | 0.010 (0.107) | 0.116 (0.502) | 0.037 (0.077) |
| CC1/2 | 1.000 (0.958) | 0.991 (0.878) | 0.998 (0.990) |
| <I/ σ(I)> | 28.3 (4.0) | 6.6 (2.6) | 7.9 (6.3) |
| Multiplicity | 20.6 (18.6) | 6.6 (6.4) | 5.3 (5.5) |
| Wilson B (Å<sup>2</sup>) | 21.85 | 59.30 | 113.89 |
| **Refinement statistics** |  |  |  |
| Reflections used in refinement | 31044 (2149) | 13281 (2634) | 1976 (1976) |
| Reflections used for R-free | 2001 (140) | 721 (154) | 130 (130) |
| R-work | 0.1727 (0.2158) | 0.2584 (0.3596) | 0.2022 (0.2022) |
| R-free | 0.1904 (0.2400) | 0.2870 (0.4261) | 0.2618 (0.2618) |
| Number of non-hydrogen atoms | 1351 | 3597 | 1211 |
| macromolecules | 1211 | 3597 | 1211 |
| ligands | 0 | 0 | 0 |
| solvent | 140 | 0 | 0 |
| Protein residues | 157 | 466 | 157 |
| RMS(bonds) (Å) | 0.008 | 0.005 | 0.003 |
| RMS(angles) (°) | 1.11 | 0.76 | 0.71 |
| Ramachandran favored (%) | 98.69 | 98.68 | 98.69 |
| Ramachandran allowed (%) | 1.31 | 1.32 | 1.31 |
| Ramachandran outliers (%) | 0.00 | 0.00 | 0.00 |
| Rotamer outliers (%) | 0.00 | 0.00 | 0.00 |
| Clashscore | 1.24 | 2.24 | 0.83 |
| Average B-factor (Å<sup>2</sup>)<sup>^</sup> | 28.87 | 105.15 | 183.05 |
| macromolecules | 27.58 | 105.15 | 183.05 |
| solvent | 40.01 | - |  |

Values in parentheses refer to the highest resolution shell. <sup>+</sup>R<sub>merge</sub> = $\sum |I - \langle I \rangle| / \sum \langle I \rangle$ , where *I* is the intensity of each individual reflection. <sup>\*</sup>R<sub>pim</sub> indicates all *I*<sup>+</sup> & *I*<sup>-</sup>. $\frac{1}{N} \sum |R_{work}| = \frac{\sum_h |F_o - F_c|}{\sum_h |F_o|}$ , where *F<sub>o</sub>* and *F<sub>c</sub>* are the observed and calculated structure-factor amplitudes for each reflection *h*. R<sub>free</sub> was determined by randomly selecting 10% of the diffraction data and excluded from refinement. Average B (Å<sup>2</sup>)<sup>^</sup> was calculated using B<sub>average</sub>.

**Table S3.** Thermodynamic parameters for the interaction of mouse Cavin1 HR1 mutants with nanobodies by ITC.

| Nanobody<br>NbB7 | K<sub>d</sub> (μM) | ΔH (kcal/mol) | ΔG (kcal/mol) | -TΔS (kcal/mol) |
| --- | --- | --- | --- | --- |
| mC1 HR1<br>Q69A |  | No binding detected |  |  |
| mC1 HR1<br>L72E |  | No binding detected |  |  |
| mC1 HR1<br>Q76A | 2.89 ± 1.01 | -1.02 ± 0.34 | -7.58 ± 0.21 | -6.55 ± 0.13 |

**Table S4.** Thermodynamic parameters for the interaction of mouse Cavin1 HR1 HT/II and TS/DD mutants with nanobodies by ITC.

| | $K_d$ (μM) | $\Delta H$ (kcal/mol) | $\Delta G$ (kcal/mol) | $-T\Delta S$ (kcal/mol) |
| --- | --- | --- | --- | --- |
| **Nanobody NbB7** |  |  |  |  |
| mC1 HR1 HT/II mutant | $0.14 \pm 0.04$ | $-4.46 \pm 0.66$ | $-9.36 \pm 0.16$ | $-4.90 \pm 0.82$ |
| mC1 HR1 TS/DD mutant | $0.067 \pm 0.02$ | $-5.44 \pm 0.37$ | $-9.82 \pm 0.22$ | $-4.37 \pm 0.58$ |
| **Nanobody NbA12** |  |  |  |  |
| mC1 HR1 HT/II mutant | $3.02 \pm 1.94$ | $-1.34 \pm 0.49$ | $-7.60 \pm 0.41$ | $-6.27 \pm 0.08$ |
| mC1 HR1 TS/DD mutant | $2.17 \pm 0.02$ | $-1.11 \pm 0.33$ | $-7.73 \pm 0.01$ | $-6.62 \pm 0.33$ |

**Table S5.** DNA constructs of cavin proteins and nanobodies.

| Species | Construct | Affinity tag | Antibiotics | Vector |
| --- | --- | --- | --- | --- |
| Mouse | Cavin1 HR1<br>(residues 45-155) | His-Ub | Amp | pHUE |
| Mouse | Cavin1 HR1<br>(residues 45-103) | His-Ub | Amp | pHUE |
| Mouse | Cavin1 HR1 5Q<br>(residues 45-155) | His-Ub | Amp | pHUE |
| Mouse | Cavin2 HR1<br>(residues 47-157) | His-Ub | Amp | pHUE |
| Zebrafish | Cavin1a HR1<br>(residues 73-193) | His-Ub | Amp, Cam | pHUE |
| Zebrafish | Cavin4a HR1<br>(residues 16-123) | His-Ub | Amp, Cam | pHUE |
| Zebrafish | Cavin4b HR1<br>(residues 12-133) | His-Ub | Amp, Cam | pHUE |
| Mouse | Cavin1 HR1 Q69A<br>(residues 45-155) | His-Ub | Amp | pHUE |
| Mouse | Cavin1 HR1 L72E<br>(residues 45-155) | His-Ub | Amp | pHUE |
| Mouse | Cavin1 HR1 Q76A<br>(residues 45-155) | His-Ub | Amp | pHUE |
| Mouse | Cavin1 full-length<br>(residues 1-392) | His-MBP | Amp | 2K-T |
| Mouse | Cavin1 HR1<br>(residues 45-155) | GST | Kan, Cam | pGEX-4T2 |
| Mouse | Cavin1 HR1 I102-I105 (HT/II) | GST | Kan, Cam | pGEX-4T2 |
| Mouse | Cavin1 HR1 D105-D106 (TS/DD) | GST | Kan, Cam | pGEX-4T2 |
| Mouse | Cavin1 Q69A<br>(residues 1-392) | GFP | Kan | pEGFP-N1 |
| Mouse | Cavin1 L72E<br>(residues 1-392) | GFP | Kan | pEGFP-N1 |
| Mouse | Cavin1 Q76A<br>(residues 1-392) | GFP | Kan | pEGFP-N1 |
| Alpaca | Nanobody NbA12 | GFP | Kan | pEGFP-N1 |
| Alpaca | Nanobody NbB7 | GFP | Kan | pEGFP-N1 |
| Mouse | Cavin1 HT/II | GFP | Kan | pEGFP-N1 |
| Mouse | Cavin1 TS/DD | GFP | Kan | pEGFP-N1 |

*\*Amp-Ampicillin; Kan-Kanamycin; Cam-chloramphenicol.*

KEY RESOURCES.

KEY RESOURCES
| REAGENT OR RESOURCE | SOURCE OR REFERENCE | IDENTIFIER |
| --- | --- | --- |
| **Bacterial Strains** |  |  |
| *E. coli* DH5α | Invitrogen | 18265017 |
| *E. coli* BL21 (DE3) | Invitrogen | C600003 |
| *E. coli* BL21 CodonPlus™ (DE3) | Agilent Technologies | 230245 |
| **Chemicals** |  |  |
| Isopropyl β-D-1-thiogalactopyranoside | Bioline | BIO-37036 |
| Benzamidine hydrochloride hydrate | Sigma Aldrich | B6506 |
| Deoxyribonuclease I (DNase I) | Sigma Aldrich | DN25 |
| Imidazole | Sigma Aldrich | 792527 |
| TALON® resin | Clontech | 635503 |
| Glutathione Sepharose 4B | GE Healthcare | GEHE17-0756-0 |
| Folch fraction I | Sigma Aldrich | B1502 |
| **Software** |  |  |
| XDS | (Kabsch, 2010) | http://xds.mpimf-heidelberg.mpg.de/ |
| AIMLESS | (Evans and Murshudov, 2013) | http://www.ccp4.ac.uk/html/aimless.html |
| Phaser | (McCoy et al., 2007) | http://www.phaser.cimr.cam.ac.uk/index.php/Phaser_Crystallographic_Software |
| Phenix | (Adams et al., 2010) | https://www.phenix-online.org/ |
| Coot | (Emsley and Cowtan, 2004) | https://www2.mrc-lmb.cam.ac.uk/personal/pemsley/coot/ |
| Pymol | (Schrodinger, USA) | https://pymol.org/2/ |
| **Other items** |  |  |
| HiLoad™ 16/600 Superdex™ 200 | GE Healthcare | 10223894 |

